# Lysosome-related organelle genes are required for mitochondrial transformations during bacterial infections in *Caenorhabditis elegans*

**DOI:** 10.64898/2026.08.04.742833

**Authors:** Jennifer D. Cohen, Ken Nguyen, David H. Hall, Frederick M. Ausubel, Gary Ruvkun

## Abstract

Detecting and responding to pathogenic bacteria is an essential function of eukaryotic cells. As bacteria-derived organelles, mitochondria carry lipids, proteins, and other molecules such as iron which are required for bacterial growth, and thus are subject to pathogenic bacterial attack. In fact, mitochondrial morphology and function are often altered during early bacterial infection. However, the mechanism by which bacterial pathogen attack triggers mitochondrial responses is unknown. Here, we demonstrate that infection by the pathogenic bacterium *Staphylococcus aureus* or *Pseudomonas aeruginosa* or hypoxia leads to remodeling of the mitochondrial network in the host *Caenorhabditis elegans*. Our analysis discovers that lysosome-related organelle (LRO) genes are required for this remodeling and indicates that LROs also precipitate downstream infection-response gene expression.

## INTRODUCTION

Eukaryotes are bombarded with bacterial pathogens, and have evolved pathways to sense and respond to infection. As antibiotic effectiveness wanes, bacterial infections may respond to modulation of these eukaryotic host immune defenses (Hancock et al., 2012). Mitochondria, as descendants of bacteria, may concentrate protein and small molecular assemblies, such as iron-sulfur (FeS) clusters, that are particularly attractive to their microbial cousins. Because mitochondria contain molecules scavenged by bacterial pathogens, they have evolved to respond to bacterial attack (Tiku et al., 2020). For example, in mammalian cell culture infected with particular bacterial species, mitochondria undergo fission, separating into small units that may be degraded via autophagocytosis (Kapetanovic et al., 2026). Mitochondria can also exhibit changes in cristae structure or loss of the outer membrane during a microbial attack (Zhou et al., 2023; Li et al., 2022). Despite the frequent observation of mitochondrial remodeling in cell culture, the extent to which mitochondrial changes occur in intact multicellular organisms, how they arise, and how they relate to immune defenses remains unclear.

Many bacteria that infect humans are also pathogenic to the nematode *Caenorhabditis elegans*, allowing us to marshal the efficiencies of low-cost laboratory cultivation to study mitochondrial response to infection (Tse-Kang et al., 2025). For example, the Gram-positive pathogen *Staphylococcus aureus*, which infects humans and causes mitochondrial rounding in mammalian cell culture (Zhou et al., 2023), also infects *C. elegans* cultivated on petri dishes containing nutrient agar that allows *S. aureus* to grow (Sifri et al., 2003). *C. elegans* feeds on these bacteria, allowing bacteria to encounter the intestine, a 32-cell tube with a lumen lined by microvilli and a thick glycocalyx that at least superficially resembles that of the mammalian intestine (Dimov and Maduro, 2019). As *C. elegans* feeds on *S. aureus* growing on the surface of nutrient agar in laboratory petri dishes, *S. aureus* accumulates inside the intestinal tube and eventually degrades microvilli to consume *C. elegans* from the inside (Irazoqui et al., 2010). To counter the infection, *C. elegans* initiates a set of MAPK signaling pathways that induce sets of innate immune genes (Pukkila-Worley et al., 2012). Although many of these gene expression pathways are well mapped, the mitochondrial or vesicular remodeling that may trigger their expression remain largely unknown.

Here, we use *C. elegans* infection with a derived clinical isolate of *S. aureus*, SH1000 (O’Neill 2010), to discover dramatic mitochondrial remodeling during very early stages of infection. During early infection, we also found surprisingly dramatic shifts in the prevalence of a specific vesicle, likely to be lysosome-related organelles (LROs), an acidic vesicle previously associated with secretion and storage of immune and mitochondria-related molecules (Bowman et al., 2019; Meng et al., 2014). Examining mutants that are defective in the production of LROs demonstrated that LRO biosynthesis genes promote the induction of host innate immune genes. We also show that mitochondrial morphology is altered in conditions of hypoxia similarly to infection with *S. aureus*, and this remodeling also requires LROs. Together, these data suggest that LRO biosynthesis genes are required for mitochondrial remodeling and promote innate immune gene expression under physiological and infection stress.

## RESULTS

### Mitochondrial morphology is altered in early infection

Transmission electron microscopy (TEM) of *wild-type* (*WT*) *C. elegans* in the early stages of infection, in which *S. aureus* has not yet breached the gut lumen, revealed dramatic mitochondrial remodeling. Pathogenic bacteria can infect *C. elegans* when they travel through the tubular intestine, which contains a lumen lined by microvilli (Figure 1A). In *C. elegans* which have been fed a standard laboratory diet of non-pathogenic *Escherichia coli* OP50, tubular mitochondria are concentrated near the intestinal lumen, immediately basal to the microvilli (Figure 1B). In contrast, the intestinal mitochondria of young adult *C. elegans* feeding on a diet of *S. aureus* for 4 hours are larger, farther from the intestinal lumen, and exhibit more disorganized, circular cristae structures compared to those in *C. elegans* feeding on *E. coli* (Figure 1B-C, S1). Crucially, all experiments reported here involved early stages of infection with *S. aureus*, under the mildly pathogenic conditions of 20°C temperature and moderately nutritive NGM-agar plates, in which *S. aureus* remained extracellular, and the lumen and microvilli were intact (Figure 1B, S2). We conclude that changes to mitochondrial morphology are a feature of early infection not prompted by disruption of the luminal barrier.

**Figure 1.**
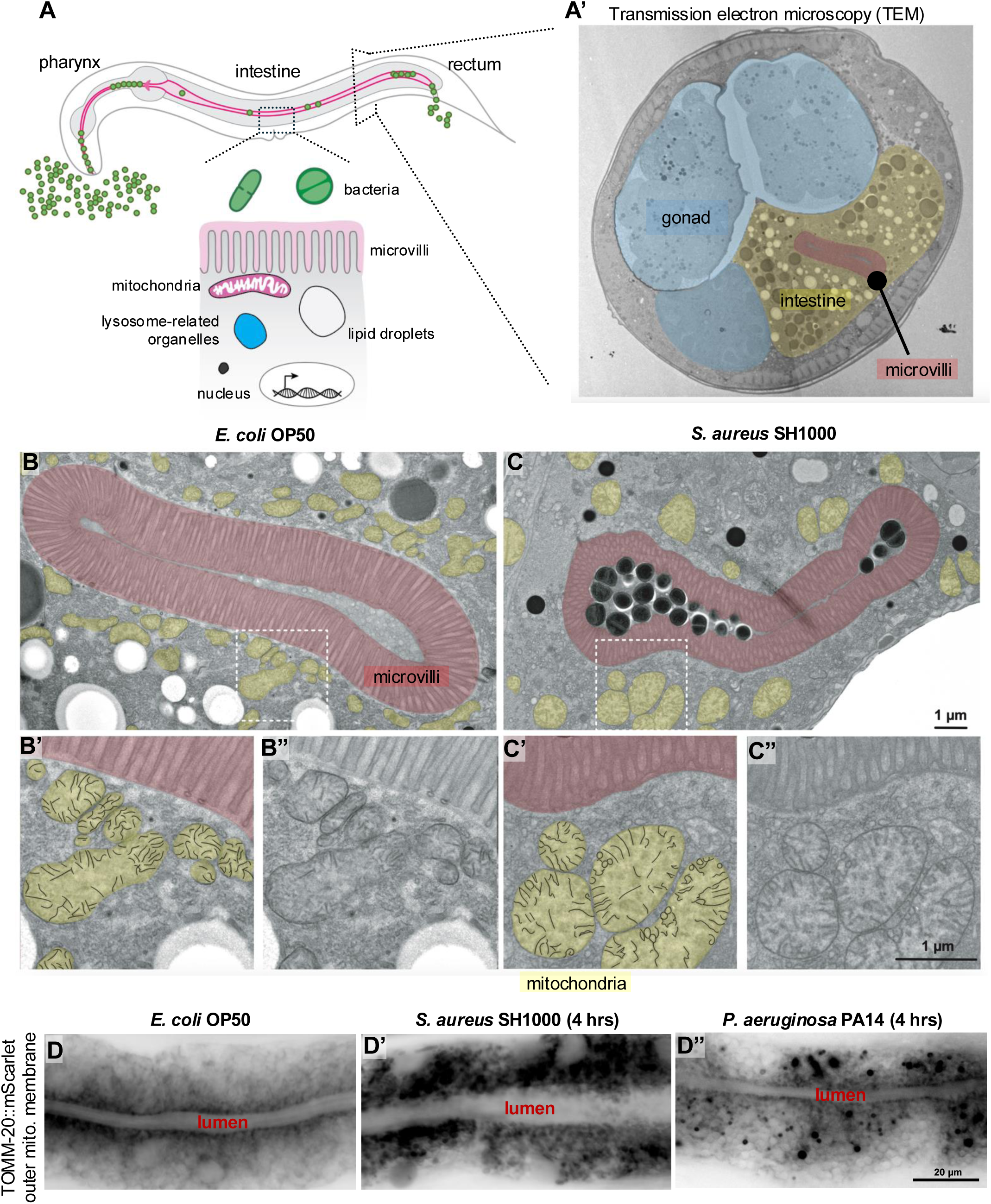
Mitochondria undergo dramatic remodeling during infection with *S. aureus* in the *C. elegans* intestine. A) Diagram of the *C. elegans* intestine. Bacteria are ingested by *C. elegans* through the pharynx, transit into the intestine, and are ejected through the rectum. While inside the intestinal lumen, bacteria pass over intestinal cell microvilli, which line the lumen. Intestinal cell cytoplasm contains a variety of organelles, including mitochondria, lysosome-related organelles (LROs), and lipid droplets. A’) TEM of *WT C. elegans*, in cross-section. Blue, gonad; yellow, intestinal cell cytoplasm; red, intestinal cell microvilli. B-B”) TEM of *WT C. elegans* intestinal cell lumen during feeding of non-pathogenic *E. coli* OP50 bacteria. Yellow, mitochondria; red, microvilli. B’-B”) Zoom-in of region highlighted in white box in B. B’) Cristae are outlined in black within mitochondria. Cristae generally have a linear appearance. B”) The same region as B’, without false coloring. C-C”) TEM of *WT C. elegans* intestinal cell lumen fed *S. aureus* SH1000. Intact *S. aureus* cells are visible within the microvilli-lined lumen. C’-C”) Zoom-in of region highlighted in white box in C. C’) Cristae are outlined in black within mitochondria. Cristae appear rounded. C”) The same region as C’, without false coloring. D-D”) Compound microscopy of *WT C. elegans* intestine expressing *vha-1pro*::TOMM-20::mScarlet, an outer mitochondrial (mito.) membrane protein, feeding on *E. coli* OP50, D’) *S. aureus* SH1000, or D”) *P. aeruginosa* PA14. The intestinal lumen is visible as a horizontal channel in which no mitochondria are present. In *C. elegans* fed *E. coli* OP50, mitochondria are clustered near the lumen. In *C. elegans* fed *S. aureus* or *P. aeruginosa* for 4 hours, mitochondria are found with disorganized or punctate localization. N > 30 for each group. All experiments were conducted on young adults selected 16 hours post mid-L4 stage.

To confirm the aberrant mitochondrial phenotype caused by *S. aureus* ingestion, we examined *C. elegans* expressing a tagged mitochondrial outer membrane protein, TOMM-20 (Mao et al., 2019). When feeding on *E. coli* OP50, TOMM-20::mScarlet signal was non-punctate and enriched along the length of the intestinal lumen (Figure 1D). However, when *C. elegans* consumed *S. aureus* for 4 hours, a time period too short to cause lethality or luminal disruption under our experimental conditions (Figure S2), TOMM-20::mScarlet was visible in larger puncta and distributed throughout the intestinal cell body, away from the lumen (Figure 1D’). We confirmed that these phenotypes required active infection by feeding *C. elegans* boiled, dead *S. aureus*, and saw no changes to mitochondrial organization: mitochondria remained enriched near the lumen, with only rare puncta (Figure S3). Roughly similar phenotypes have been observed in *C. elegans* lacking the mitochondrial Rho-GTPase gene *miro-1,* possibly due to an inability to fuse mitochondria (Mao et al., 2019), but these phenotypes have not been previously reported in the case of a pathogenic infection. These results indicate that mitochondrial morphology is widely altered in an *S. aureus* bacterial infection prior to observable damage to the intestinal lumen barrier.

To test if this mitochondria morphology phenotype was unique to infection by the Gram-positive bacterium *S. aureus*, we examined TOMM-20::mScarlet signal during infection by the Gram-negative pathogen *Pseudomonas aeruginosa* PA14. Unlike *S. aureus*, which degrades the intestinal lumen and eventually consumes the *C. elegans* host from the inside, *P. aeruginosa* PA14 is pathogenic to *C. elegans* in part via secreted toxins, including ExoA, which target translational machinery (McEwan et al., 2012; Kirienko et al., 2014). We observed similar, though less dramatic, loss of TOMM-20::mScarlet from the intestinal lumen and redistribution into puncta within the *C. elegans* intestine during early infection (4 hours) by *P. aeruginosa* (Figure 1D). We conclude that changes in mitochondrial organization are not specific to infection by *S. aureus,* but may represent a general response to cellular stress caused by bacterial infection.

As changes to mitochondrial morphology and loss of cristae structure are both linked to decreased mitochondrial function, we tested if mitochondrial stress occurred during *S. aureus* infection (Cogliati et al., 2013). Two transcriptional reporters are activated during multiple types of mitochondrial stress: *hsp-6pro*::GFP, which contains the promoter of the *C. elegans* HSP70 homolog and is upregulated during the mitochondrial unfolded protein response, and *cyp-14A4pro*::GFP, which contains the promoter of a cytochrome P450 detoxification gene that is upregulated during *P. aeruginosa* PA14 infection (Yoneda et al., 2004; Mao et al., 2019). Neither mitochondrial stress reporter was activated during the early stages of an *S. aureus* infection, despite the mitochondrial morphology differences observed (Figure 2A). Because these reporters can fail to express GFP under conditions of high sterols, which are found in *S. aureus*, we tested for additional signs of altered mitochondrial function (Marshall and Wilmoth, 1981; Liu et al., 2014). Staining with TMRM, a red dye whose fluorescence intensity is an indicator of mitochondrial membrane electron gradient, showed consistent, albeit small, increases in fluorescence intensity during infection with *S. aureus*, which implies increased mitochondrial membrane potential (Figure 2B) (Chen 1988). TMRM fluorescence intensity decreased when *C. elegans* were fed *P. aeruginosa PA14*, suggesting different mechanisms of infection for each bacterium (Figure 2B). We also observed increased fluorescence from photoconvertible genetically encoded fluorescent proteins, which indicates increased ROS production (Figure 2C) (Dooley et al., 2004; Belousov et al., 2006). Together, these data suggest mild but inconsistent changes in mitochondrial function, with slightly increased mitochondrial membrane potential. It remains possible that the changes in mitochondrial morphology observed during infection may not dramatically alter mitochondrial function.

**Figure 2.**
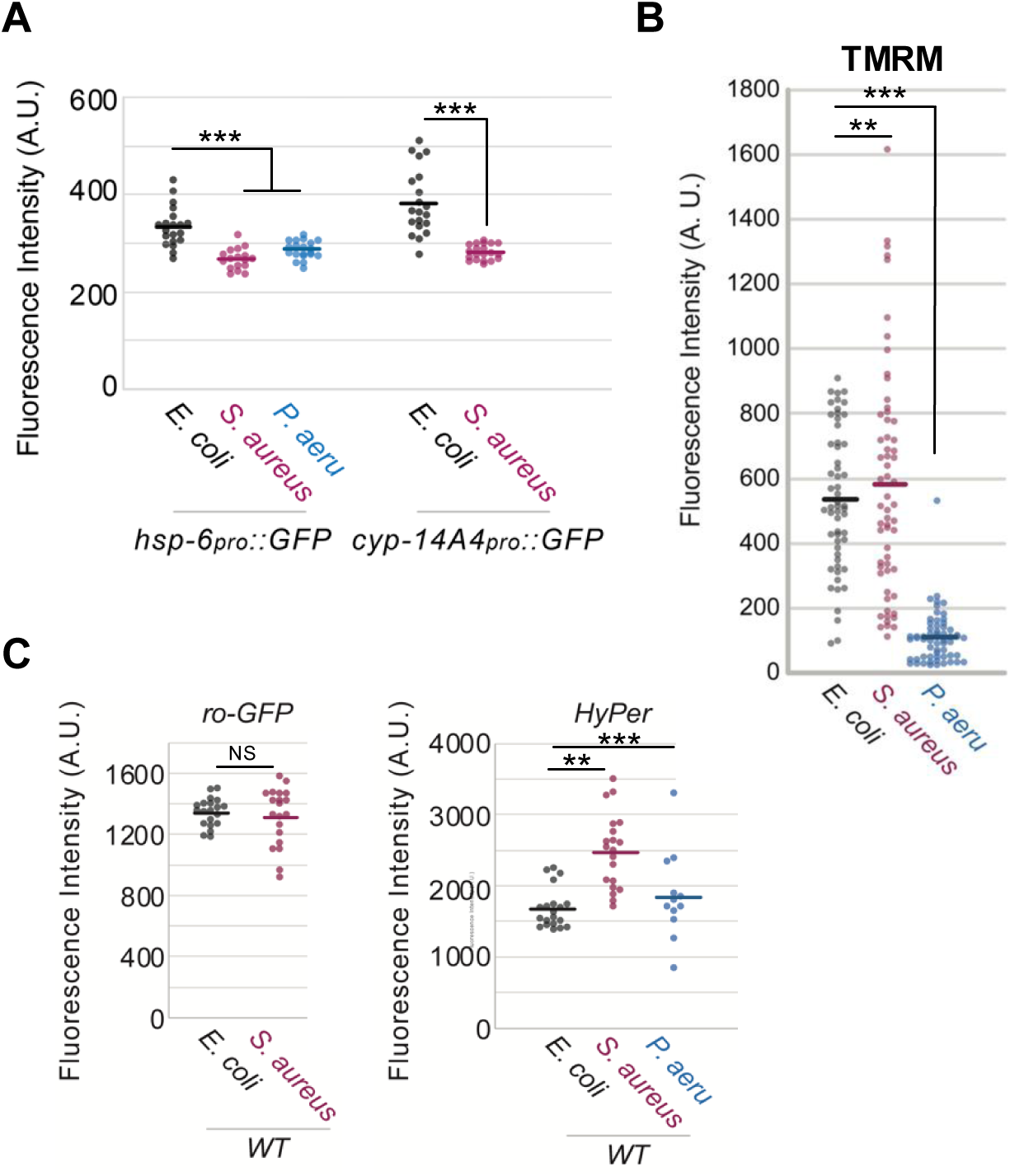
Mitochondrial function shows small shifts during early infection. A) Fluorescence expression of mitochondrial stress reporters is decreased during early infection. B) TMRM dye shows slightly increased intensity after 4 hours of *S. aureus* infection. C) Fluorescence intensity of cytoplasmic ROS reporters show inconsistent results during early infection. Ro-GFP2 detects glutathione redux potential, while HyPer detects hydrogen peroxide (Back et al., 2012). Fluorescence intensity was measured by averaging (mean gray value) the fluorescence intensity of two 48 µm circular regions of interest (ROIs) in the anterior portion of the intestine. Dots represent individual *C. elegans*.

### Mitochondria are found near autofluorescent LRO contents during infection

To more closely identify the contents of mitochondrial puncta observed during *S. aureus* infection, we examined *C. elegans* containing endogenously-tagged inner and outer mitochondrial membrane proteins (SCPL-4 and TOMM-70, respectively) (Figure 3; Figure S4) (Valera-Alberni et al., 2024). In *C. elegans* consuming non-pathogenic *E. coli* OP50, mitochondria line the intestinal lumen, with rare blebs near the basal surface of intestinal cells. But after only 4 hours consuming the bacterial pathogens *P. aeruginosa* or *S. aureus*, mitochondria were less dense along the intestinal lumen and showed increased punctate localization of mitochondrial proteins (Figure 3A). Despite the formation of puncta, the fluorescence intensity of each protein remained steady or decreased during early infection, suggesting that mitochondrial number dropped overall (Figure S4). Importantly, we also observed that mitochondrial puncta tended to co-localize with a blue auto-fluorescent signal that is produced by a kynurenine pathway intermediate and stored in LROs (Coburn et al., 2013; Buceta et al., 2019; Del Borello et al., 2019; O’Rourke and Ruvkun, 2013; Clokey and Jacobson 1986). We confirmed the association of mitochondria with LROs by imaging mitochondria with GLO-1::GFP, a Rab32/38 protein required for LRO synthesis (Figure 3D) (Morris et al., 2018). These data confirm altered mitochondrial localization and indicate some degree of colocalization between mitochondria and LROs.

**Figure 3.**
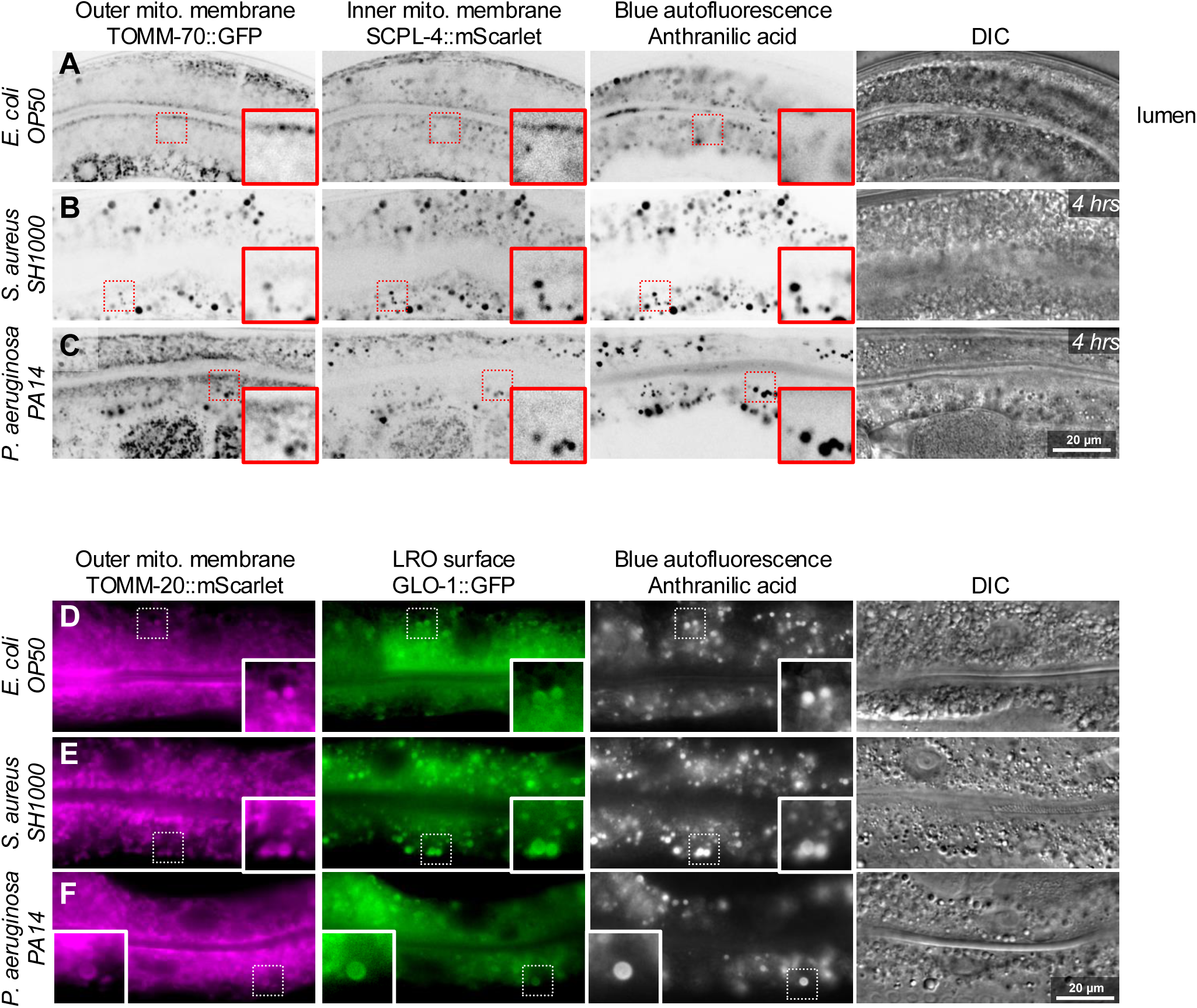
Dysmorphic mitochondria co-localize with LROs. A-C) Confocal microscopy of young adult *WT C. elegans* intestines fed *E. coli OP50* or transferred to *S. aureus* or *P. aeruginosa* for 4 hours. A) Both outer and inner mitochondrial membrane proteins are enriched near the luminal surface, visible as a narrow line, between which no signal is present. B-C) This luminal-enriched localization is lost in *C. elegans* fed *S. aureus* or *P. aeruginosa* for 4 hours. 10 µm x 10µm insets (red box) show puncta which co-localize with blue autofluorescent material (anthranilic acid) housed in LROs. These puncta are more common in *C. elegans* fed *S. aureus* and *P. aeruginosa* for 4 hours. D-F) Compound microscopy of young adult *C. elegans* expressing mgTi33 [*vha-6pro*::*TOMM-20::mScarlet*] and kxEx15 [*glo-1pro::GLO-1::GFP*]. Rare mitochondrial puncta are visible in *C. elegans* fed *E. coli OP50*, but puncta become common in *C. elegans* fed *S. aureus* or *P. aeruginosa* for 4 hours. Puncta frequently co-localize with GLO-1::GFP, both known to localize to LROs. n >30 for all groups.

### LROs may be lost during infection

Alongside mitochondrial remodeling, we also observed differences in vesicle density by TEM in *S. aureus* infected *C. elegans* (Figure 4A). The *C. elegans* intestine contains many vesicles, including lipid droplets, lysosomes, LROs, and yolk granules (Dimov and Maduro, 2019). To identify which vesicles were perturbed in *C. elegans* consuming *S. aureus*, we classified vesicles by electron density. Pale, low electron density vesicles, which may be lipid droplets, underwent a slight drop in frequency during infection that did not reach statistical significance (Figure 4B) (Zhang et al., 2010). In contrast, moderately electron dense vesicles, previously hypothesized to be LROs, showed a dramatic drop in frequency during *S. aureus* infection (Figure 4B’) (Zhang et al., 2010). Lastly, rare, very electron dense vesicles of unknown etiology underwent a moderate increase in frequency during infection (Figure 4B”). Close inspection revealed that these latter vesicles were not bound by a cell envelope and thus are unlikely to represent engulfed *S. aureus* (Figure S1B). We conclude that only one major class of vesicle is missing during *S. aureus* infection, and hypothesize that this vesicle class is LRO.

**Figure 4.**
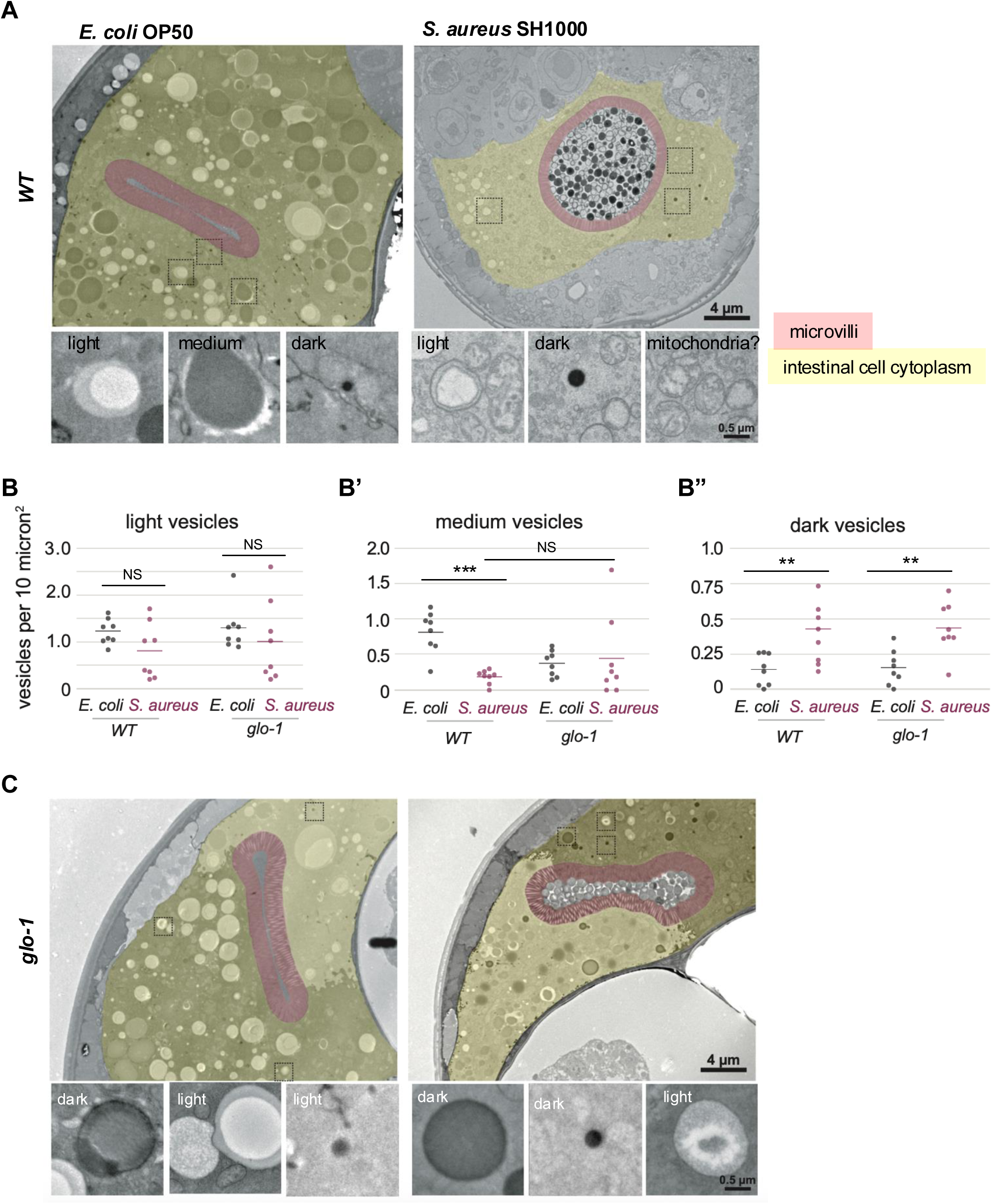
LRO-like vesicles are lost during early infection with *S. aureus*. A) TEM of *WT C. elegans* intestinal cytoplasm after feeding *E. coli* OP50 or *S. aureus* SH1000. Red, microvilli; yellow, intestinal cell cytoplasm. Insets are examples of light, medium, and dark electron density vesicles magnified from images above. Medium electron density vesicles are virtually absent in *WT C. elegans* infected with *S. aureus*. B-B”) Quantification of light, medium, and dark electron density vesicles per micron^2^ of intestinal cell cytoplasm, excluding microvilli and luminal space. *** = p< 0.001,** = p < 0.01. Mann Whitney t-test. C) TEM of *glo-1* mutant *C. elegans* intestinal cytoplasm after feeding *E. coli* OP50 or *S. aureus* SH1000. Insets are examples of light, medium, and dark electron density vesicles magnified from images above. Medium electron density vesicles are virtually absent in *glo-1 C. elegans* regardless of feeding *E. coli* or *S. aureus*.

By TEM, we observed that the *C. elegans glo-1* mutant, which lacks LROs, had fewer vesicles overall in intestinal cells under non-pathogen-infection conditions (Figure 4B, 4C). By classifying vesicles by electron density as before, we observed the same frequency of both low and high electron density vesicles as in *WT C. elegans*, but a clear lack of medium density vesicles compared to *WT C. elegans* (Figure 4B-B”). Because these medium-density vesicles are nearly absent in *glo-1* mutants defective for LRO production, we propose that they represent LROs. *S. aureus* infection did not alter the number of vesicles observed by TEM in *glo-1* mutants, possibly because LROs were already absent (Figure 4B-B”). These data are consistent with a model in which the vesicles lost in *WT C. elegans* during *S. aureus* infection are LROs.

We observed rare examples of medium-electron-density vesicles contacting mitochondria in a cloud of wrinkled membranes in *S. aureu*s-infected *C. elegans*, suggesting that putative LROs may fuse with mitochondria to change their morphology (Figure S5A). LRO-like vesicles were observed near mitochondria in non-infected *C. elegans*, but never appeared to share membranes (Figure S5B). While these experiments are not sufficient to rule out alternative mechanisms for LROs acting on mitochondria, fusion between LROs and mitochondria during infection would be consistent with the continued presence of LRO contents (GLO-1::GFP, blue autofluorescence), but the loss of distinct LRO-like vesicles as imaged by TEM.

### Mutants lacking LROs do not develop mitochondrial morphology remodeling in infection

Because fluorescent microscopy revealed partial colocalization of mitochondrial puncta and LROs, and TEM revealed that LROs may vanish during *S. aureus* infection, we hypothesized that LROs act on mitochondria during infection to promote mitochondrial morphology remodeling. We therefore examined several *C. elegans* mutants that are unable to synthesize LROs. These mutants were originally identified by the lack of autofluorescent material in the intestine, but were later identified as homologs of highly conserved genes (Hermann et al., 2005; de Voer et al., 2008). Loss of these genes causes Hermansky-Pudlak syndrome in humans, characterized by defects in secretion of melanin and clotting factors as well as immunodeficiency (Wang and Young 2025). *glo-1*, a homolog of human Rab32/38, localizes to LROs and is required for their synthesis. *glo-4* encodes a guanine nucleotide exchange factor that acts on GLO-1 and is a homolog of human HERC4 (de Voer et al., 2008). *pgp-2* encodes an ABC transporter of unknown function. These mutants are slightly short and wide (Dumpy; Dpy) but have a normal lifespan and do not express stress response pathways under baseline conditions (Figure S6A) (Hermann et al., 2005; Morris et al., 2018; Schroeder et al., 2007; Hajdú et al., 2023; Tan et al., 2024). These mutants show decreased expression of some stress responsive genes relative to *WT* after exposure to a chemical irritant (Hajdú et al., 2023). LROs were also reported to contain the immune regulator TIR-1, and thus are implicated in the infection response (Tse-Kang et al., 2024). LROs are synthesized by a conserved pathway and are involved in infection and stress responses in both humans and *C. elegans,* though the mechanism by which LROs sense and respond to stress remain unknown.

To test if mitochondria respond to *S. aureus* in *C. elegans* lacking LROs, we examined *glo-1* mutants. TEM revealed that in *glo-1* mutants, normal tubular mitochondrial morphology is retained both on non-pathogenic *E. coli* OP50 and on *S. aureus*. *glo-1* mutant mitochondria also maintained normal cristae structure when grown on *E. coli* (Figure 5A). Tracings of individual mitochondria from TEM micrographs demonstrated that mitochondria of *WT* and *glo-1 C. elegans* became larger in both area and perimeter during infection with *S. aureus* (Figure 5B). In *WT C. elegans*, the area increased more than the perimeter, indicating that mitochondria became more rounded during infection (Figure 5C). In contrast to *WT C. elegans*, the mitochondria of the *glo-1* mutant became larger evenly, with an unchanged area to perimeter ratio, indicating that mitochondria retained their tubular shape during infection (Figure 5C). In *WT C. elegans* fed *E. coli* OP50, many mitochondria appeared in direct contact with the base of microvilli at the intestinal lumen. Fewer mitochondria had this localization in *WT C. elegans* fed *S. aureus*. Again, *glo-1* mutant mitochondria showed a less dramatic difference, with approximately equal numbers of mitochondria at the base of microvilli during infection (Figure 5D). Importantly, *glo-1* mutants developed an enlarged intestinal lumen and slowed defecation while consuming *S. aureus* to a similar degree as *WT C. elegans*, demonstrating similar levels of infection (Figure 6A, S6B). Together, these results indicate that during *S. aureus* infection of *WT C. elegans*, mitochondria become larger and more rounded, and migrate away from the lumen, but *glo-1* mutant mitochondria fail to round and retain their luminal localization.

**Figure 5.**
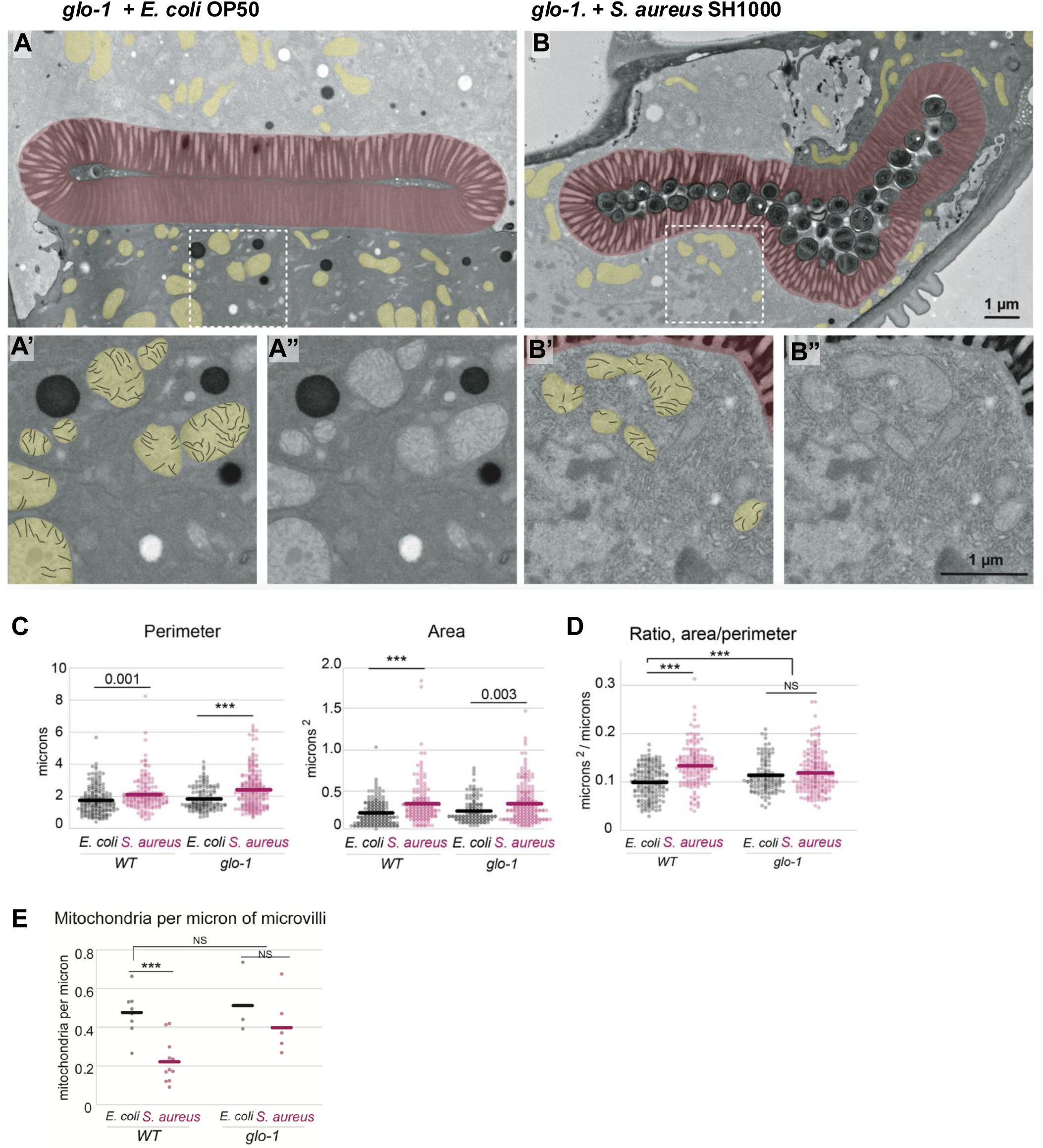
LRO mutants show decreased mitochondrial remodeling during *S. aureus* infection. A-A”) TEM of *glo-1* mutant *C. elegans* intestinal cell lumen during feeding on non-pathogenic *E. coli* OP50 bacteria. Yellow, mitochondria; red, microvilli. A’-A”) Zoom-in of region highlighted in white box in A. A’) Cristae are outlined in black within mitochondria. Cristae generally have a linear appearance. A”) The same region as A’, without false coloring. B-B”) TEM of *glo-1 C. elegans* intestinal cell lumen fed *S. aureus* SH1000. Intact *S. aureus* cells are visible within the microvilli-lined lumen. B’-B”) Zoom-in of region highlighted in white box in B. B’) Cristae are outlined in black within mitochondria. Cristae appear linear. B”) The same region as B’, without false coloring. C) ImageJ-generated measurements of mitochondria from manual tracings of 10 unique TEM micrographs per group. D) The ratio of area to perimeter from measurements depicted in C. E) Number of mitochondria touching the base of microvilli per micron of luminal length as measured manually from TEM images. Mann Whitney t-test. *** = p < 0.001.

**Figure 6.**
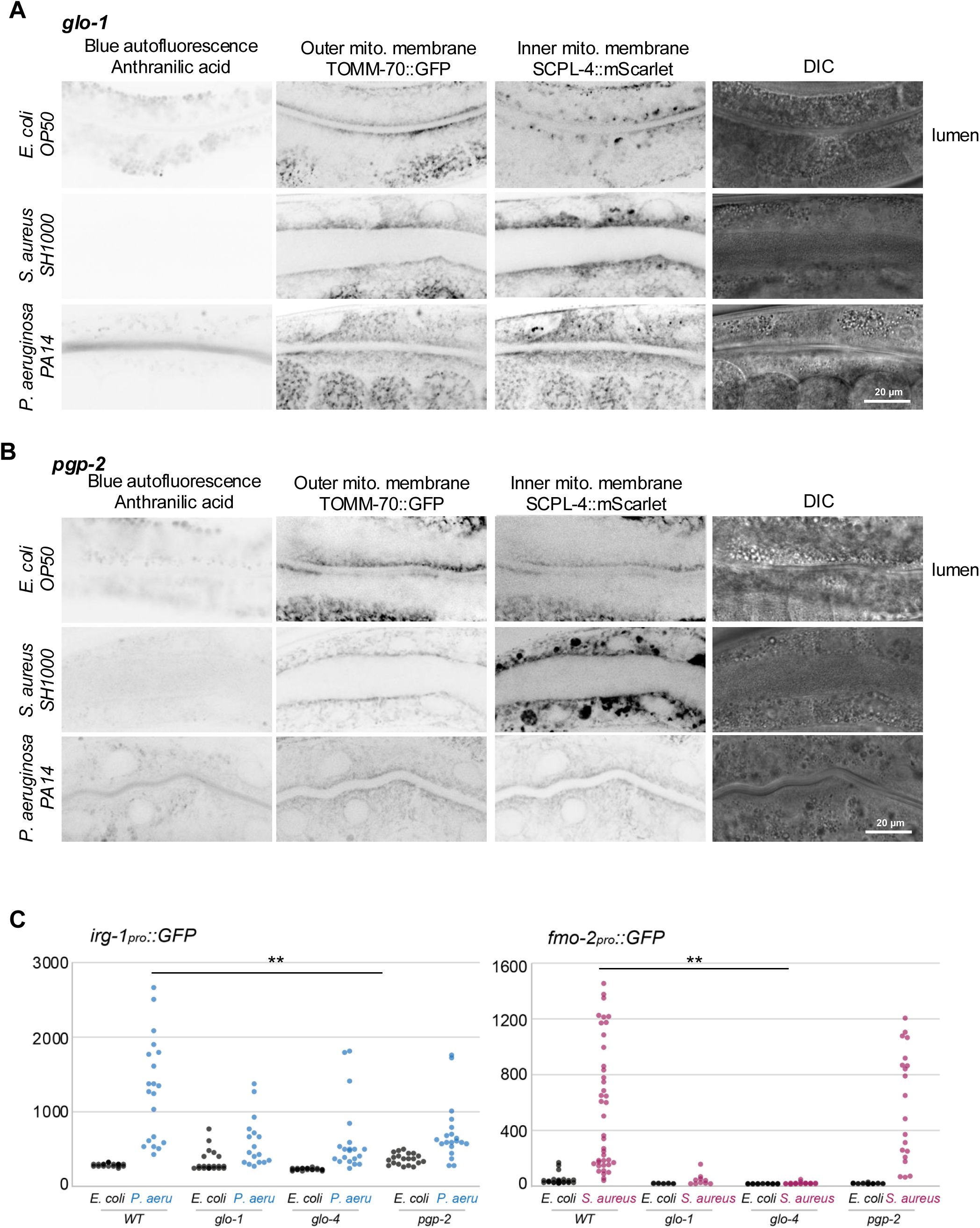
LRO genes promote mitochondrial remodeling. A-B) Confocal microscopy of young adult *WT C. elegans* intestines fed *E. coli* OP50 or transferred to *S. aureus* or *P. aeruginosa* for 4 hours. Both outer and inner mitochondrial membrane proteins are enriched near the luminal surface, visible as a narrow line, between which no signal is present. This luminal-enriched localization is maintained in *glo-1* and *pgp-2 C. elegans* fed *S. aureus* or *P. aeruginosa* for 4 hours. Blue autofluorescent material is not visible in *glo-1* and *pgp-2* mutants as these mutants do not synthesize the LROs (Hermann et al., 2005). C) Quantification of intestinal fluorescence from transgenes that are expressed during infection by *P. aeruginosa* (*irg-1pro*::GFP) or *S. aureus* (*fmo-2pro*::GFP). At infection timespans less than 16 hours, *glo-1* and *glo-4* mutants show muted fluorescence expression.

We confirmed normal mitochondrial morphology during *S. aureus* infection in *glo-1 C. elegans* using the endogenous fluorescently-tagged mitochondrial membrane reporters used previously (TOMM-20::mScarlet, TOMM-70::mScarlet, SCPL-4::GFP). The punctate mitochondrial pattern observed in *WT* during *S. aureus* infection was not observed in *glo-1*, *glo-4*, or *pgp-2* mutants (Figure 3, 6). Although the intestinal lumen became distended during early *S. aureus* infection, mitochondria continued to localize close to the lumen (Figure 5E, 6A, B). We conclude that LRO genes are required for mitochondrial remodeling during infection.

### LRO genes are required for full expression of immune genes

We next investigated the role of LRO-induced mitochondrial remodeling in the infection response. Infection by *S. aureus* or *P. aeruginosa* elicit specific gene expression programs that promote immunity (Wang and Young 2025). Loss of *glo-1* and *glo-4* mutants moderately decreased the expression of the reporter genes (*fmo-2pro::GFP, irg-1pro::GFP*) normally responsive to each bacterial pathogen (Figure 6C) (Leiser et al., 2015; Estes et al., 2010). We conclude that Glo mutants promote full expression of stress-responsive genes.

To test if *glo-1* mutants were more susceptible to infection than *WT*, we examined behavioral and survival phenotypes. Although some previous reports showed that *glo-1* mutants were slightly more susceptible to bacterial pathogens and chemical stressors than *WT*, we did not observe significant differences in lifespans for *glo-1* mutants vs. *WT* on non-pathogenic *E. coli* OP50 and pathogenic *S. aureus* (Figure S6A) (Hajdú et al., 2023). Similarly, *glo-1* mutants slowed their rate of defecation, an indicator of infection, when feeding on *S. aureus* to the same degree as *WT* (Figure S6B) (Rae et al., 2012). However, we did observe substantially increased embryonic and larval arrest in *glo-1* mutants vs. *WT* raised on *S. aureus* strains (Figure S6C). Therefore, *glo-1* mutants had dramatically more *WT*-appearing mitochondria during infection, and the organismal impact was observed in offspring of *C. elegans* raised on *S. aureus*.

### Mitochondria also respond to hypoxia via LRO genes

One way that infection could be sensed by *C. elegans* is the shortage of specific nutrients, which produces cellular stress. Mitochondrial electron transport typically requires oxygen and heme, both of which can be consumed by *S. aureus*. To test if mitochondria display similar morphological remodeling during stress to those seen in *S. aureus* infection, we examined hypoxic conditions. We observed that *WT C. elegans* mitochondria remodel during hypoxia in a manner superficially similar to their remodeling during *S. aureus* infection (Figure 7A, 1C). In contrast, *glo-1* and *pgp-2* mutants showed muted or absent changes in mitochondrial morphology during hypoxia, even when also infected by *S. aureus* (Figure 7B, C). These results indicate LROs are required for mitochondrial morphology remodeling, which are a general response to many types of stress (Figure 8).

**Figure 7.**
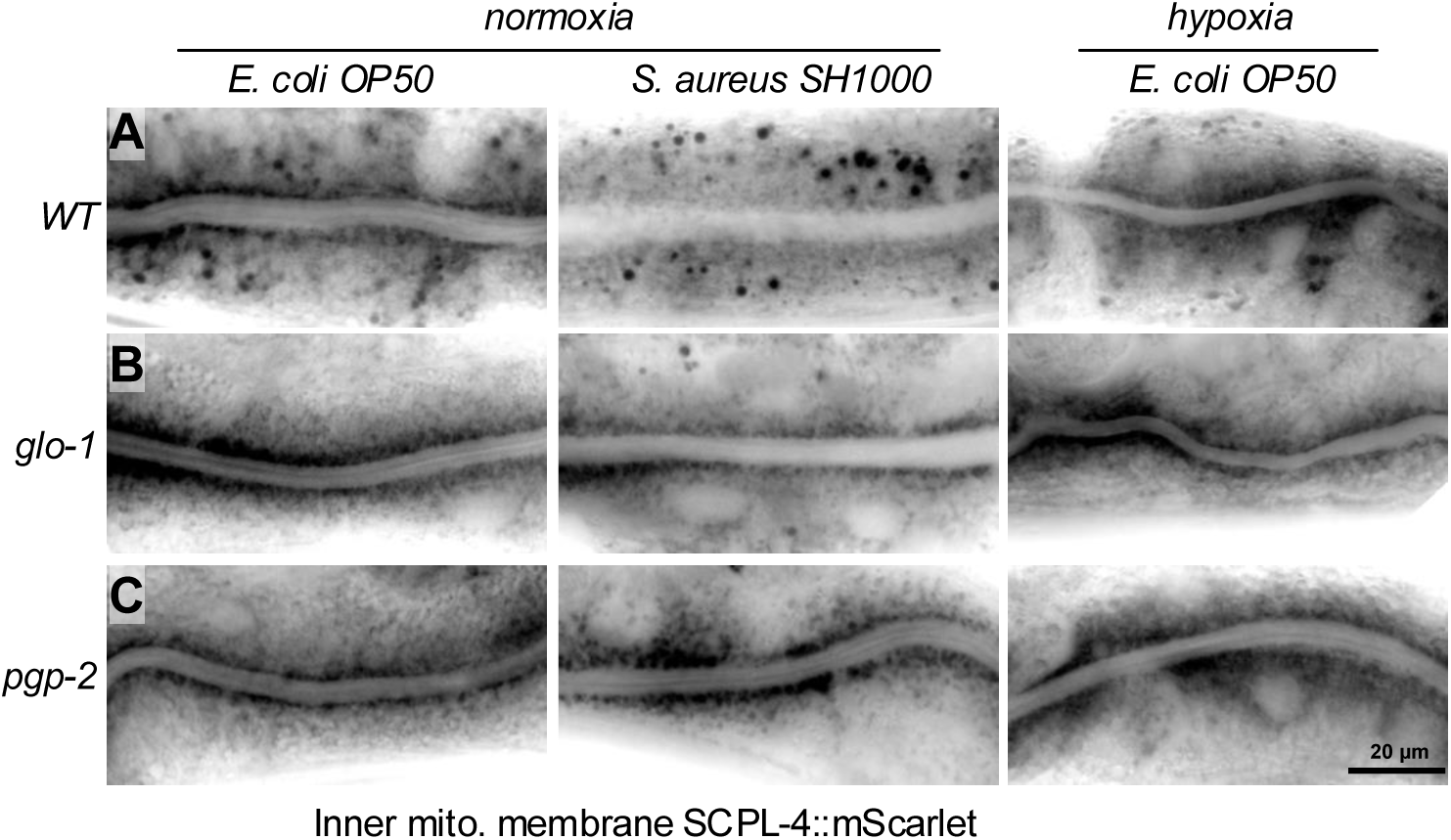
Hypoxia leads to superficially similar mitochondrial remodeling as *S. aureus* infection. Compound microscopy of *WT C. elegans* intestine expressing SCPL-4::mScarlet, an inner mitochondrial (mito.) membrane protein, from the endogenous *scpl-4* locus. The intestinal lumen is visible as a horizontal channel in which no mitochondria are present. A) In *C. elegans* fed *E. coli* OP50 under normoxia, mitochondria are clustered near the lumen, but this localization is lost in *WT C. elegans* fed *S. aureus* or placed under hypoxia (1% oxygen) for 4 hours while feeding on *E. coli*, mitochondria are found with disorganized or punctate localization. B-C) *glo-1* and *pgp-2* mutants maintain relatively normal mitochondrial localization under both *S. aureus* feeding and hypoxia. n > 20 for each group. All experiments were conducted on young adults selected 16 hours post mid-L4 stage.

**Figure 8.**
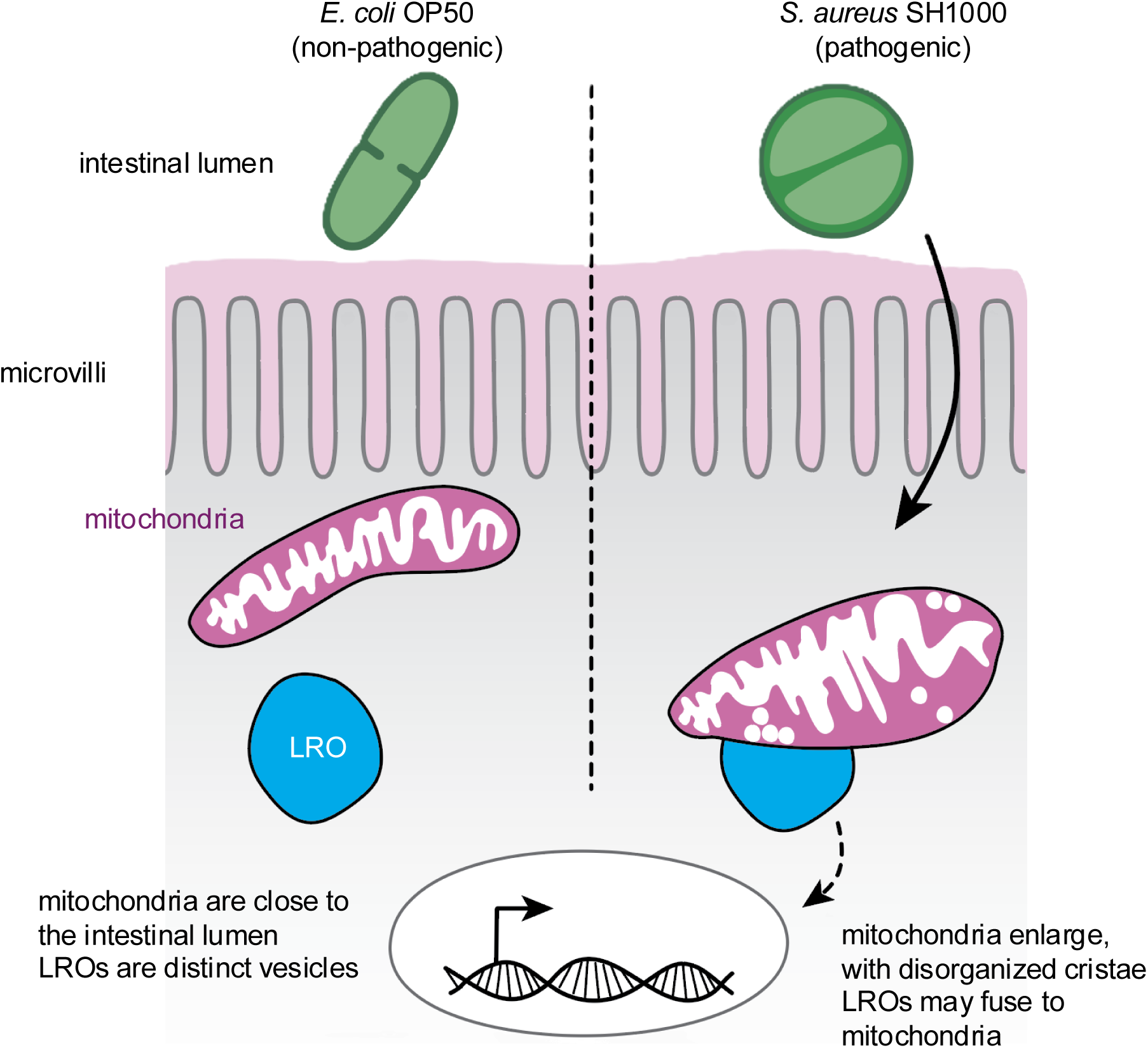
LROs promote mitochondrial remodeling during infection to trigger transcriptional immune response. Bacteria in the intestinal lumen of *C. elegans* signal to LROs via an unknown mechanism. LROs then may interact with mitochondria to change mitochondrial morphology. This sequence of events then promotes expression of innate immune response pathways.

## DISCUSSION

By investigating early stages of infection and stress responses, we discovered that the intestine responds to pathogenic bacterial infection with dramatic changes to mitochondrial morphology and loss of LRO vesicles. We propose that LRO vesicles associate with mitochondria during early infection, leading to loss of LROs and producing rounded mitochondria that no longer localize to the intestinal lumen. Mutants that fail to synthesize LROs showed no change in mitochondrial morphology and failed to express innate immune genes. We also demonstrate that mitochondrial remodeling mediated by LRO vesicle pathway genes are part of a critical step in innate immune gene activation. In all, we have discovered that LROs, an evolutionarily conserved vesicle implicated in immunity, may promote immune responses via triggering mitochondrial remodeling during infection and physiological stress.

### LROs may act as mitochondrial regulators

In humans, mutations in LRO genes cause Hermansky-Pudlak syndrome, a recessive genetic disorder that causes immunosuppression (Wang and Young 2025). Although mitochondria are not traditionally thought of as primarily involved in this syndrome, one study of lung tissue in patients indicated the presence of abnormally shaped and functionally impaired mitochondria (Cuevas-Mora et al., 2021). Cell culture studies have also shown defects in peroxisome and lipid metabolic function in this syndrome, which could derive from defects in ATP accumulation (Kook et al., 2016). Future work should elucidate the extent to which LROs support mitochondrial function, particularly during infection, in humans.

Because LROs are the site of maturation for melanosomes and promote secretion of clotting factors, they are often considered secretory vesicles (Bowman et al., 2019). However, they are also known to store mitochondrial-related compounds in *C. elegans*: zinc, heme, and blue fluorescent anthranilic acid, an intermediate of the kynurenine pathway that is a precursor to rhodoquinone and NAD+ (Coburn et al., 2013; Buceta et al., 2019; Del Borello et al., 2019; Roh et al., 2012; Chen et al., 2018). Rhodoquinone is an electron transport alternative to ubiquinone, functional in hypoxia (Salinas et al., 2020); zinc is a cofactor for mitochondrial superoxide-dismutases and can bind heme precursors in the absence of iron (Brito et al., 2026; Labbé et al., 1999); heme supports mitochondrial complex IV (Poulos 2014). In mammals, the storage sites of all three substances are unknown. Whether these substances are stored together in LROs in mammals, and how such co-storage might influence mitochondrial activity during infection, also remains unknown. Based on the data in this paper, we propose that LROs may deliver stored mitochondrial cofactors to stressed mitochondria under conditions of cellular stress or hypoxia created by bacterial infection.

Though there are many reports of mitochondrial fission or altered morphology in response to bacterial infection in mammalian cell culture, the function of this remodeling remains unknown. In this work, we have linked this remodeling to interaction with LROs and downstream expression of immune genes. However, it remains unclear to what extent mitochondrial function is compromised. Modest results with TMRM dye suggest no consistent change in mitochondrial membrane potential during infection with *S. aureus*, and it is possible that instead of compromising function, the mitochondria of *C. elegans* simply shift to using rhodoquinone, which regenerates NAD+ from NADH in the absence of oxygen, rather than using the oxygen-requiring ubiquinone to complex III and IV pathways (Salinas et al., 2020).

Previous work in *C. elegans* has implicated LROs in immunity. In *C. elegans*, LRO synthesis is triggered in part by HLH-30, a TFEB homolog (O’Rourke and Ruvkun, 2013). HLH-30 is in turn required for expression of several innate immune genes during *S. aureus* infection (Visvikis et al., 2014). These findings suggest that the innate immune role for HLH-30 may rest primarily in promoting the synthesis of fully developed LROs. How HLH-30 promotes LRO synthesis, and whether it also impacts loading of LROs with rhodoquinone or other cargo, remains unknown. The immune regulator TIR-1, a homolog of mammalian SARM1 that activates the conserved p38 PMK-1 mitogen-activated protein kinase (MAPK) host defense pathway, is also localized to the membrane of LROs and its aggregation is inhibited in intact LROs (Liberati et al., 2004; Tse-Kang et al., 2024; Tse-Kang and Pukkila-Worley, 2024). It remains to be tested whether interaction between LROs and mitochondria might trigger altered activity of TIR-1.

### How do LROs associate with mitochondria, and by what trigger?

Our data suggest that LROs interact with mitochondria during bacterial pathogen infection. Although there are many reports of mitochondrial fission during infection or cellular stress, and many more of contact points between mitochondria and vesicles like the ER or lysosomes, this is the first report of contact between mitochondria and LROs during infection. We further demonstrate that this interaction, whether direct or indirect, is functional: in the absence of LROs, mitochondrial morphology remains normal and innate immune genes are less robustly expressed. However, it remains unclear if LROs bind to the mitochondrial surface, and what molecules enable or trigger the interaction.

This work demonstrated similar phenotypes during the early stages of infection by two very different bacterial pathogens and physiological stress: *S. aureus* is a Gram-positive, opportunistic pathogen that expresses no known toxins and instead apparently infects by degrading microvilli on the intestinal luminal surface; *P. aeruginosa* is a Gram-negative, aggressive pathogen that expresses translational elongation-specific toxins which contribute to killing *C. elegans;* and hypoxia is a specific limitation in oxygen availability, with no infection. It remains unclear how these very different stresses are detected and how that detection triggers LRO interaction with mitochondria. Because these bacterial pathogens trigger expression of distinct sets of genes, LROs must represent only one of several innate immune pathways activated by bacterial infection. However, the fact that both these very different bacteria trigger a common response – LRO interaction with mitochondria – suggests that there may be an undiscovered “master switch” to indicate infection. Future work should test more bacterial pathogens and the sole known *C. elegans* virus, Orsay, to discover how broadly such a “master switch” might act.

## MATERIALS and METHODS

### Worm strains, alleles and transgenes

All strains were derived from Bristol N2 and were grown at 20°C under standard conditions (Brenner, 1974). *C. elegans* strain N2 *WT* was obtained from the Caenorhabditis Genetic Center (CGC). See Table S1 for a complete list of strains generated in this study.

### Microbial Strains

*E. coli* OP50, *S. aureus* SH1000, and *P. aeruginosa* PA14 were cultured overnight in LB at 30°C. 100 μl of liquid culture was seeded on NGM plates free of antibiotics. To ensure active cultures,

*C. elegans* were added to plates within 5-6 hours of seeding. Plates were stored at 20-25°C before the addition of *C. elegans* and 20°C after *C. elegans* were added.

### Infection

L4 larvae were staged by vulva lumen morphology and aged 16 hours to generate day 1 adults. Day 1 adults were transferred to freshly seeded *E. coli*, *S. aureus*, or *P. aeruginosa* from overnight cultures prepared as discussed above.

### Staging and microscopy

Live worms were immobilized with 50 mM levamisole in M9 and mounted on a slide with 2% agarose. Fluorescent, brightfield, and Differential Interference Contrast (DIC) images were captured on a Zeiss AX10 Zoom.V16 microscope fitted with a Leica DFC360 FX camera or with a Leica TCS SP8 confocal microscope (Leica, Wetzlar Germany). Images were processed and merged using ImageJ/FIJI. TMRM dye was resuspended in M9 at 10µM concentration. 10µL of TMRM dye mix was added to PCR tubes with *C. elegans* and the indicated bacteria. *C. elegans* was allowed to incubate for 45 minutes in the dye mix, then was removed to a fresh NGM agar plate seeded with the indicated bacteria. *C. elegans* was incubated on the fresh plate for 20 minutes, and then was transferred to agar pads with levamisole for imaging. Protocol adapted from (Mallick and Haynes, 2024).

For TEM, L4 hermaphrodites from N2 (*wild-type*) or JJ1271 (*glo-1*(*zu391*)) strains were transferred from *E. coli* OP50 to fresh plates containing lawns of either *E. coli* OP50 or *S. aureus* SH1000, then incubated overnight. Though this exposure was longer than that used for fluorescence experiments (4 hours), the intestinal lumens of *C. elegans* remained intact, possibly because *S. aureus* was prepared in advance on agar plates and may have been in stationary, not log phase, during experiments (Figure 1C). *C. elegans* were then fixed by high pressure freezing followed by freeze substitution into osmium tetroxide in acetone (Weimer, 2006), and then rinsed and embedded into LX112 resin and cut into serial thin sections of approximately 70 nm each. Sections were observed on a JEM-1010 transmission electron microscope (Jeol, Peabody Massachusetts). Images were processed in ImageJ and manually pseudo-colored in Adobe Illustrator (Adobe, San Jose California). We imaged intestines for n = 8 N2 fed *E. coli*, n = 8 N2 fed *S. aureus*, n = 5 *glo-1* fed *E. coli*, and n = 10 *glo-1* fed *S. aureus*. Data for all animals are shown.

### Lifespan analysis

Animals were synchronized by egg-laying and picked at L4 stage in groups of 100, recorded as day 0. Adults were separated from their progenies by manually transferring to new plates. Survival was examined on a daily basis. The average of three independent trials containing 100 *C. elegans* each is shown.

### Defecation quantification

Day 1 adult *C. elegans* were transferred onto fresh lawns of either *E. coli* OP50 or *S. aureus* SH1000 and allowed to equilibrate for 1 hour. Each individual was followed by light microscopy for a total of 3 defecations and the interval between each defecation was recorded.

### Quantification of TEM

Mitochondrial area and perimeter were measured by manual tracing and the Shape Descriptor measurement tool in ImageJ. To measure the number of mitochondria proximal to the intestinal lumen, all mitochondria touching the base of the microvilli, where an electron-dense cytoskeletal band is present, were counted. The cytoskeletal band was then manually traced and its length measured as a proxy for the perimeter of the lumen. The number of mitochondria touching the base of microvilli was divided by the length of the lumen at the base of the microvilli for mitochondria per micron. Because some worms are more darkly stained than others, vesicle electron density was only compared within individual micrographs. Vesicles were counted as light, medium, or dark based on their appearance relative to neighboring vesicles. Statistical tests performed were unpaired Student’s t-tests.

### Quantification of fluorescence microscopy

To quantify transgene fluorescence intensity, a 48-micron diameter circle was drawn and two measurements were taken per worm, at the most anterior point of the intestine and then just behind that. Both measurements were anterior to the gonad, which in some cases obstructed microscopy of the intestine. Measurements from each worm were averaged. Shown are results from 3 independent experiments, n > 10 for each experiment.

For quantification of the spatial organization of mitochondrial fluorescence in the intestine, a 10-micron thick line was drawn across the intestine, from the basal end of one side of the intestine across the lumen to the basal end of the opposing side and fluorescence intensity along the line was collected. The midpoint of the lumen was noted from DIC images and was used to align measurements from different animals. Fluorescence intensities were normalized with 1 equal to the highest value. Average and standard deviation values were reported. n > 12 for each set of measurements. All fluorescence intensities were captured using FIJI/ImageJ.

## ACKNOWLEDGEMENTS

We thank all members of the Ruvkun laboratory for their much-needed advice. Sam Wattrus offered invaluable advice and help with confocal microscopy. Ken Nguyen and David Hall performed TEM on *C. elegans*. Fred Ausubel and Gary Ruvkun provided guidance in experimental design and critical reading of the manuscript.

Support for this work comes from NIH grant awarded to 5F32GM146400 to Jennifer D. Cohen, HHMI grant GMAS Fund 279015, and NIH grants 5R01AG016636 and 5R01GM044619 awarded to Gary Ruvkun. Some strains were provided by the CGC, which is funded by NIH Office of Research Infrastructure Programs (P40 OD010440).

---

**Figure S1.**
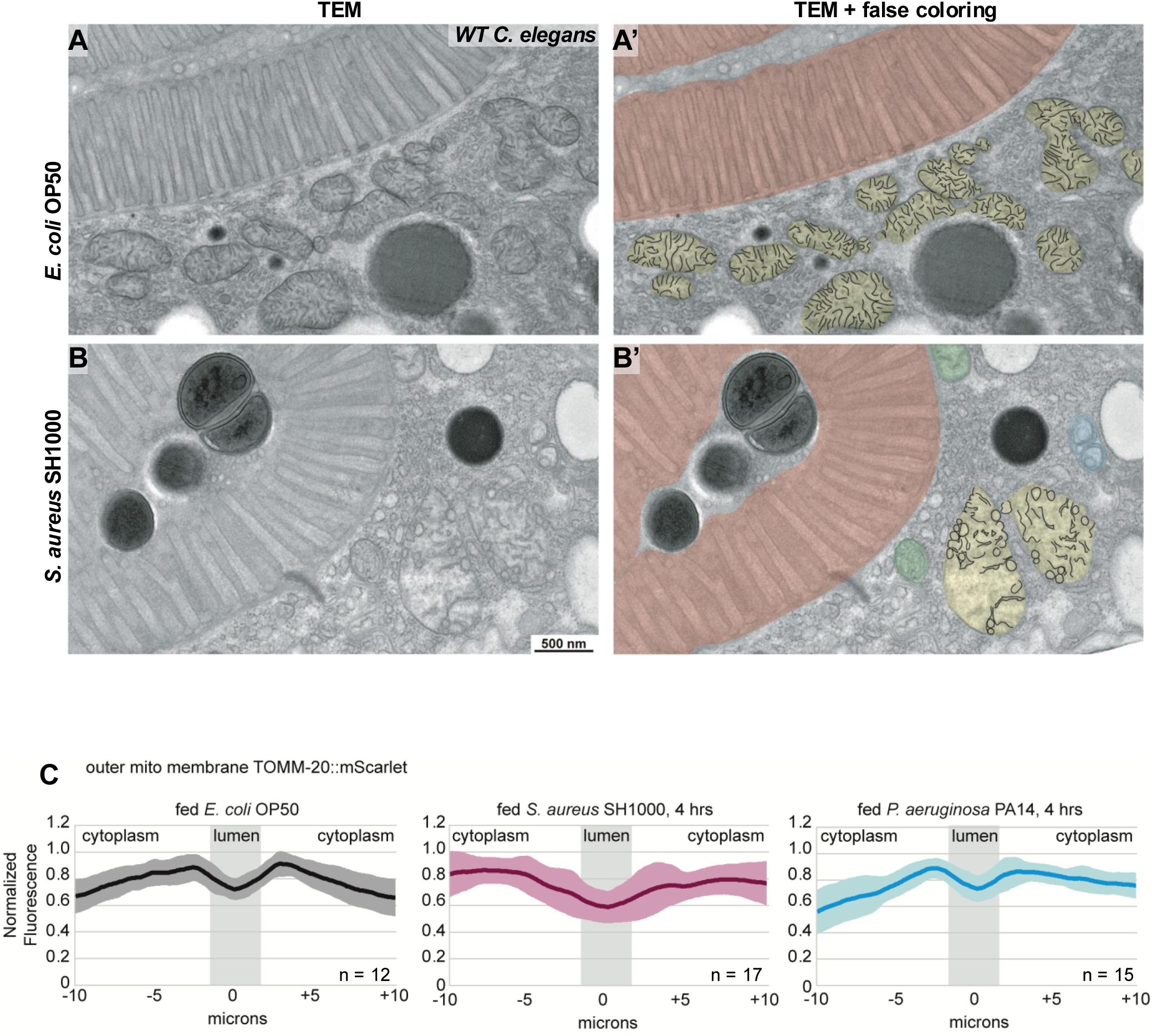
Mitochondrial cristae are disorganized during infection though *S. aureus* remains extracellular. A) TEM of *C. elegans* intestinal lumen fed *E. coli* OP50. A’) Same images as A, with false coloring. B) TEM of *C. elegans* intestinal lumen fed *S. aureus* SH1000. B’) Same image as B, with false coloring. Red, microvilli; yellow, mitochondria; blue and green, additional vesicles of unknown identity visible in *C. elegans* consuming *S. aureus*. Cristae are outlined in black. *S. aureus* cells are visible inside lumen, adjacent to microvilli. Electron-dense vesicles resemble *S. aureus* superficially, but are not encased in a cell envelope and thus are unlikely to be *S. aureus*. Mitochondria appear more rounded and with circular cristae in *C. elegans* fed *S. aureus*. Arrowheads; circular cristae seen only in *C. elegans* fed *S. aureus*. C) Quantification of TOMM-20::mScarlet signal in the intestine of *C. elegans* fed *E. coli* OP50, or transferred to *S. aureus* or *P. aeruginosa* for 4 hours. A line 10 µm thick was drawn across the intestine and average fluorescence intensity (mean gray value) was recorded. Intensity plots for each animal were normalized and centered at the midpoint of the lumen. Plots are shown for 10 µm on either side of the luminal midpoint. Averages of all animals are shown (dark line) alongside standard deviation for each point (lighter colored shading). The approximate location of the lumen, based on combined DIC images of each animal, is indicated (gray bar). Values taken from cytoplasm, where mitochondria are present, is indicated with a white background. More mitochondrial signal is present near the lumen, and less is found basal to the lumen, in *C. elegans* fed *E. coli* OP50 continuously. *C. elegans* fed *P. aeruginosa* also show some enrichment of mitochondria near the intestinal lumen, but *C. elegans* fed *S. aureus* show no enrichment of mitochondria near the lumen, with mitochondrial signal dispersed throughout the intestinal cell cytoplasm. n = 12, *E. coli* sample; n = 17, *S. aureus* sample; n = 15, *P. aeruginosa* sample.

**Figure S2.**
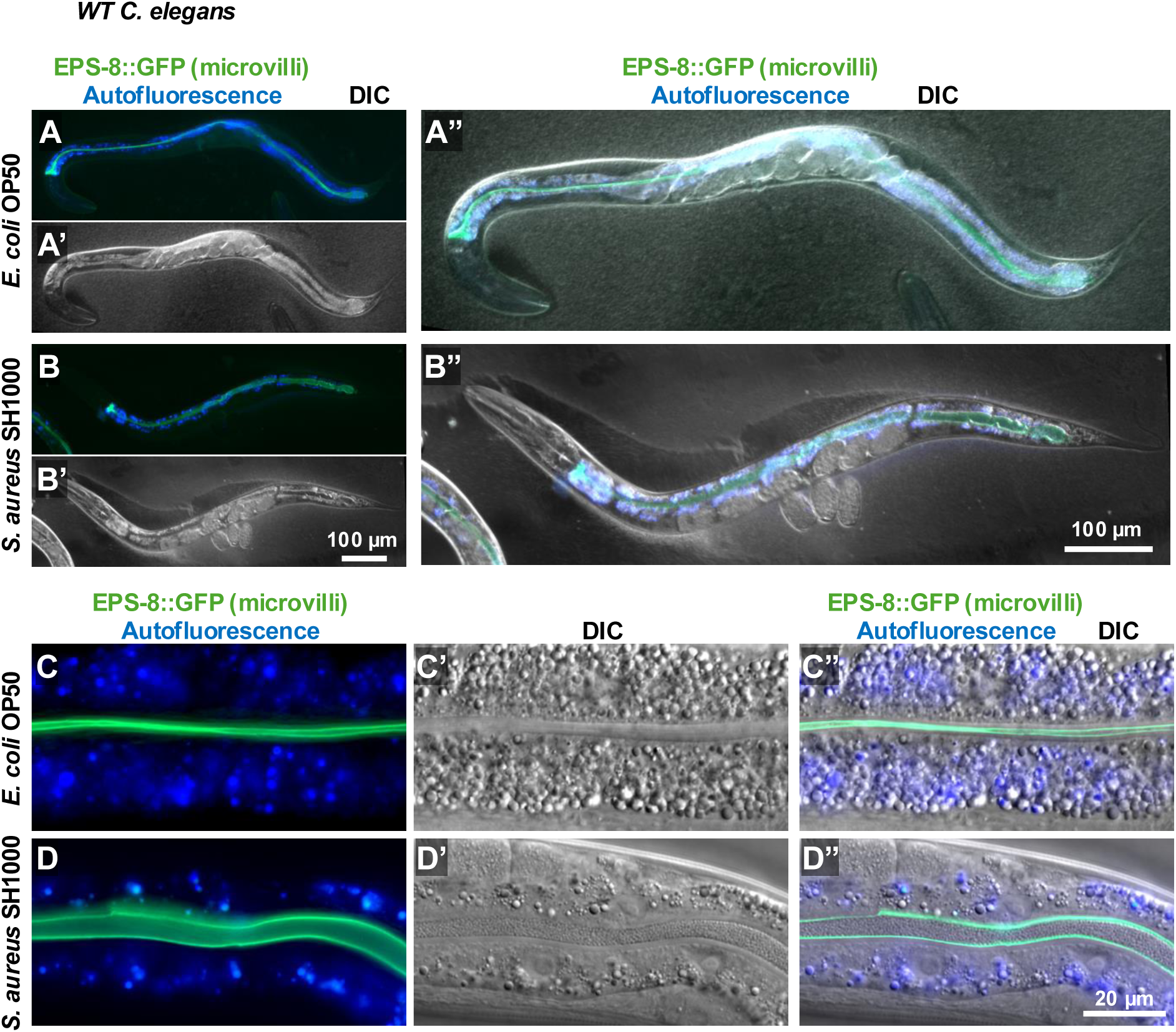
*C. elegans* retains a normal morphology during early infection with *S. aureus*. Compound microscopy of *C. elegans* expressing EPS-8::GFP from its endogenous locus. EPS-8 has actin capping activity and localizes to the outer portion of microvilli (Croce et al., 2004; Zhang et al., 2023). A-B) *C. elegans* retains normal gross morphology after 4 hours feeding on *S. aureus* SH1000. C-D) Higher magnification reveals an intact microvilli layer (labeled by EPS-8::GFP) in *C. elegans* fed *S. aureus*. D) The lumen is visibly distended (greater distance between green microvilli layers). D’-D”) *S. aureus* cells are visible as round objects within the lumen. Blue autofluorescence is present in the intestinal cytoplasm. n > 10 for each group.

**Figure S3.**
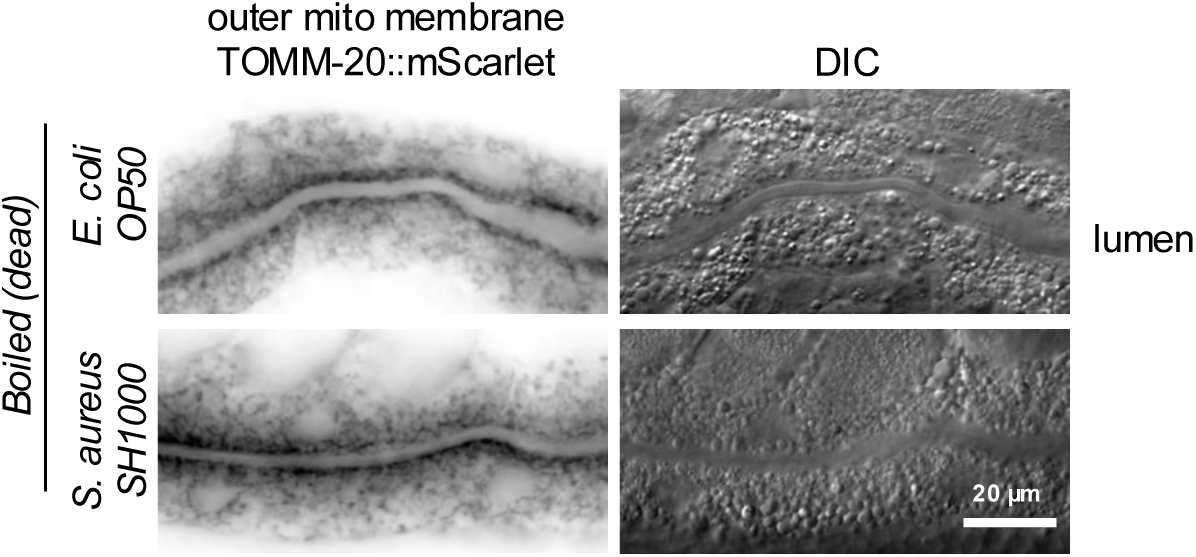
Dead *S. aureus* does not induce mitochondrial morphology remodeling. Bacterial cultures were boiled for 2 hours then plated on *C. elegans* NGM plates. Young adult *C. elegans vha-6p::tomm-20(1–54)::mScarlet* was then allowed to feed on the dead bacterial cultures for 4 hours before imaging by compound microscopy. Mitochondria were visible along the intestinal lumen as in *C. elegans* fed live *E. coli* OP50, with no punctate or disorganized localization as seen in *C. elegans* transiently fed *S. aureus* SH1000 (Figure 1D-D’). The lumen is visible as a negative space across the horizontal midline of each image, where mitochondrial signal is absent. n = 20 for each group.

**Figure S4.**
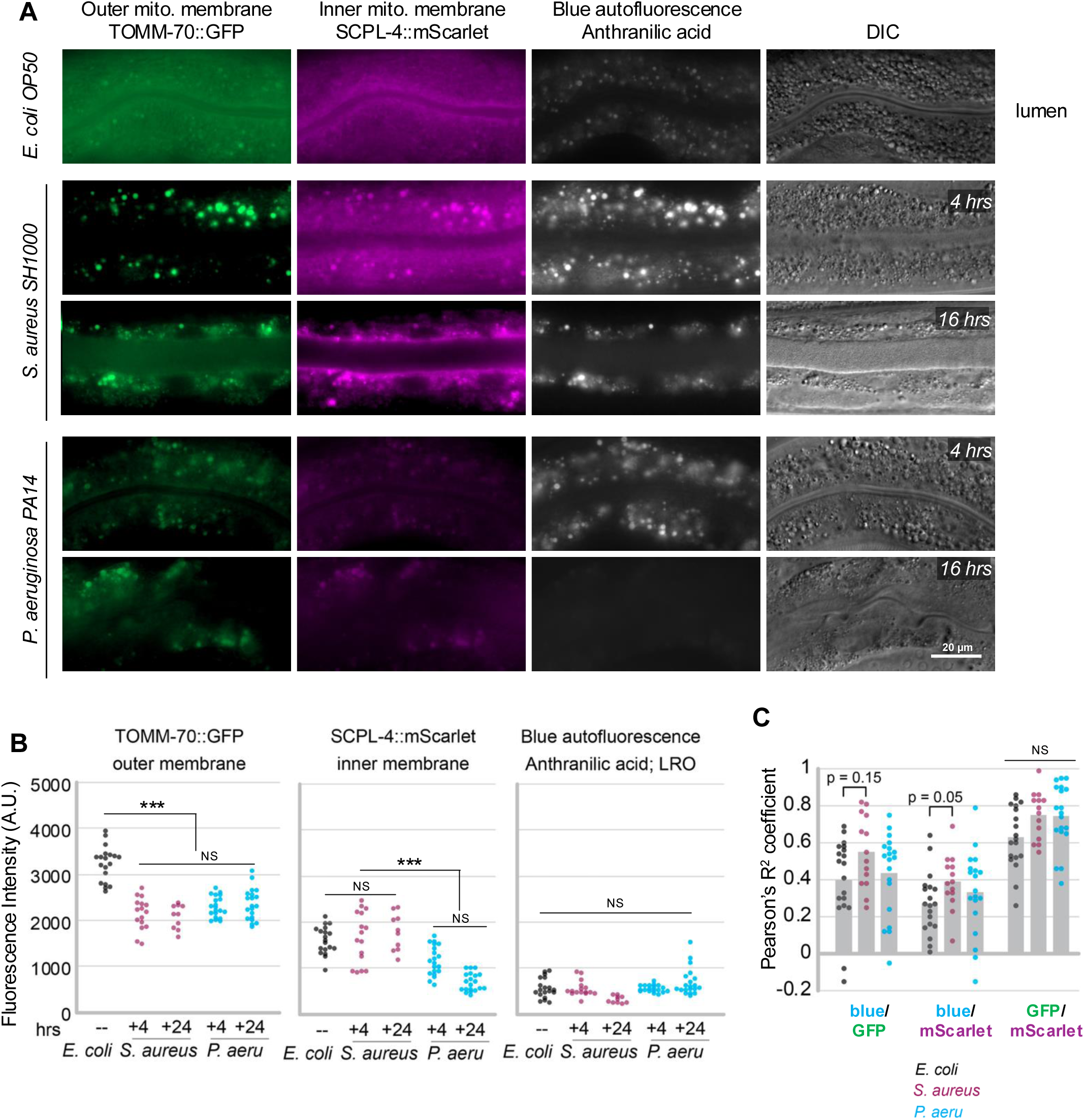
Dysmorphic mitochondria accumulate during infection. A) Compound microscopy of *WT* young adult *C. elegans* expressing endogenously-tagged TOMM-70::GFP and SCPL-4::mScarlet and fed *E. coli*, *S. aureus*, or *P. aeruginosa* for 4 or 16 hours. B) Fluorescence intensity of each channel was measured by averaging (mean gray value) the fluorescence intensity of two 48 µm circular regions of interest (ROIs) in the anterior portion of the intestine. C) Colocalization between blue autofluorescent material (housed in LROs), TOMM-70::GFP, and SCPL-4::mScarlet was performed using Pearson’s Correlation Coefficient in ImageJ. Briefly, a ROI containing the intestine was identified in each image based on presence of blue autofluorescence. Colocalization within that ROI was then automatically calculated in ImageJ.

**Figure S5.**
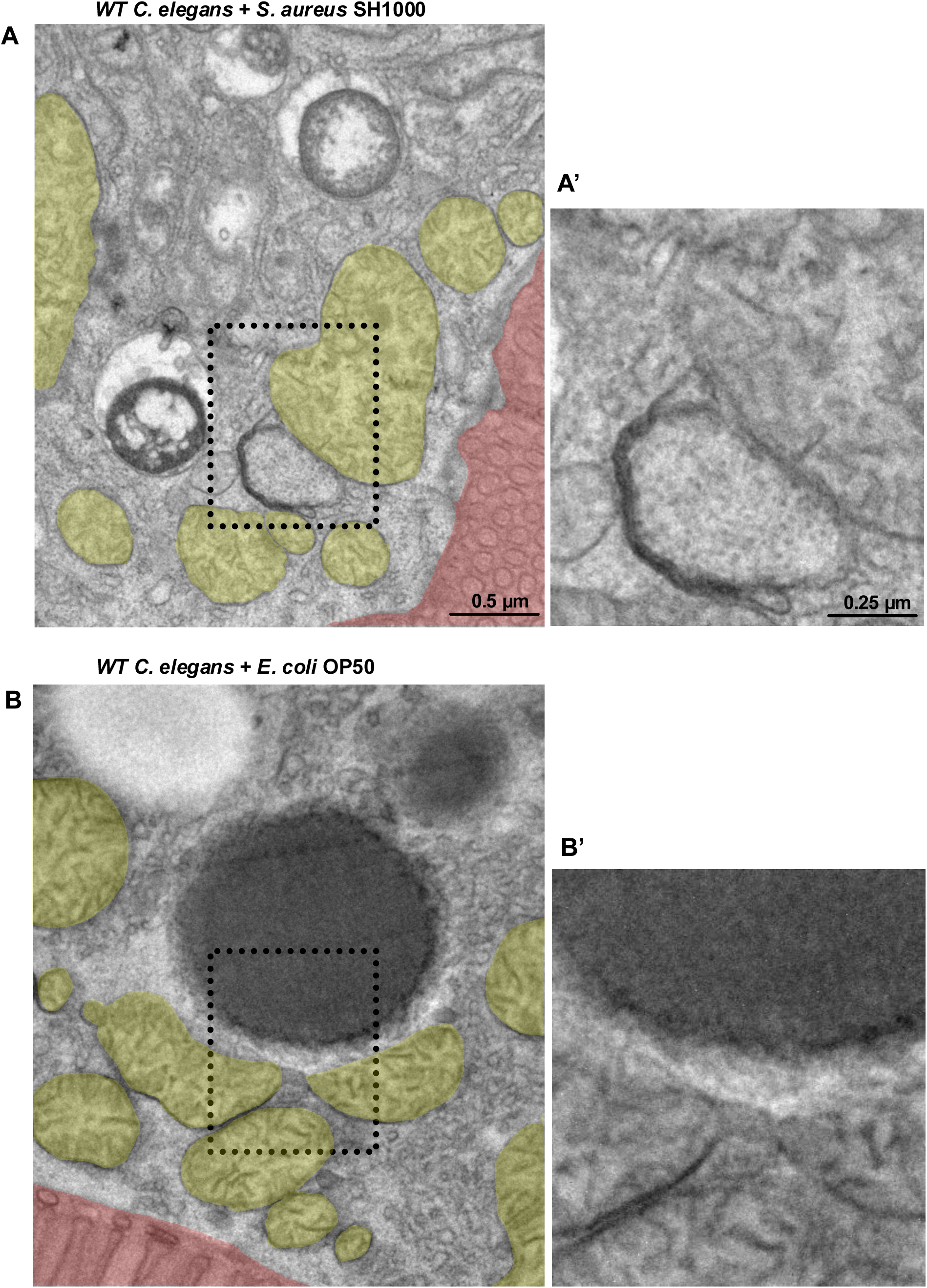
LRO-like vesicles may contact mitochondria during infection. A) TEM micrograph of the intestinal lumen. Mitochondria; yellow. Microvilli; red. A vesicle of medium electron density appears next to, and possibly continuous with the outer membrane of, a mitochondrion. A’) Zoom-in of the same image without false coloring.

**Figure S6.**
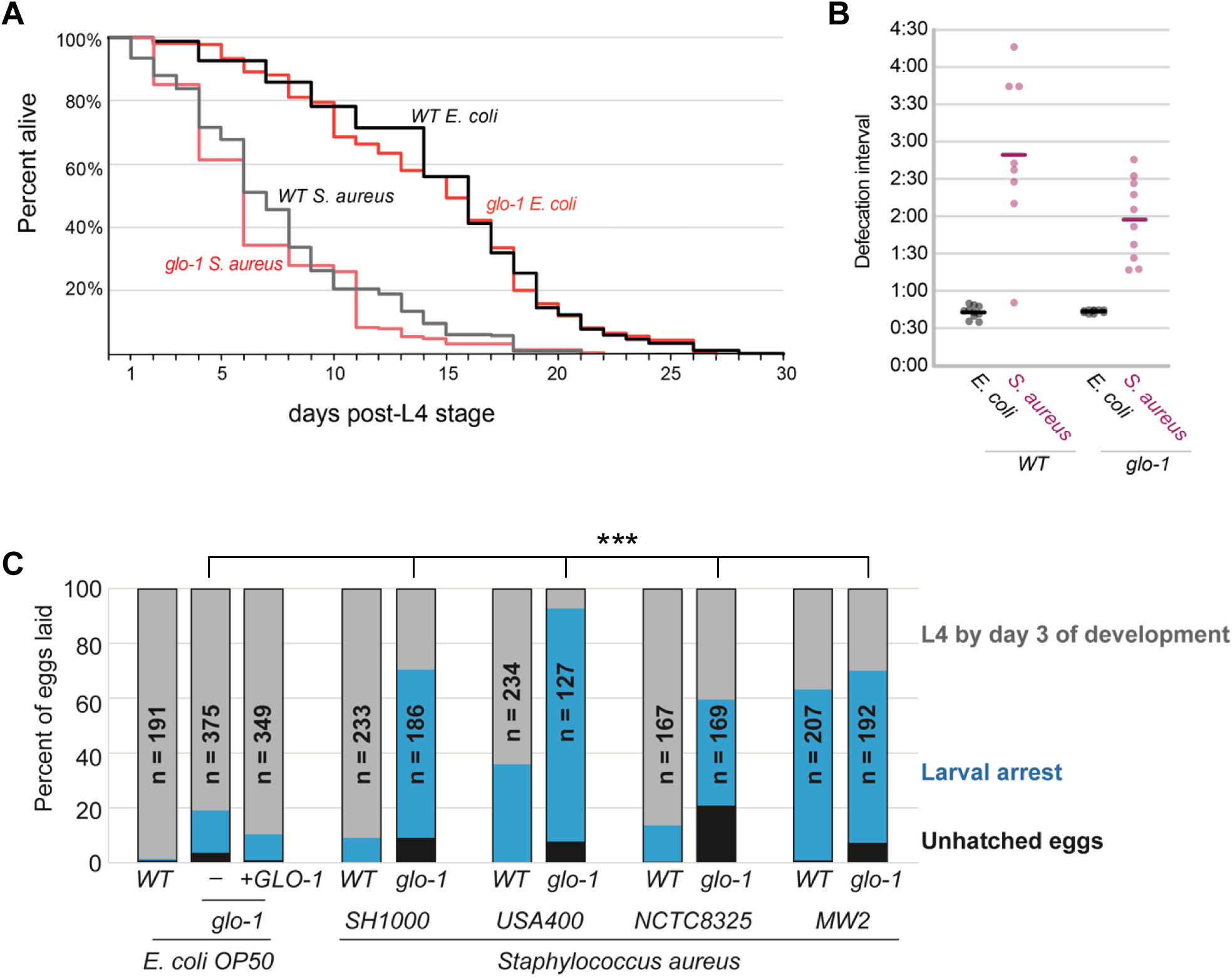
*glo-1* mutants respond like *WT* to infection with *S. aureus*. A) Lifespan quantification of *WT* and *glo-1 C. elegans* fed *S. aureus* on NGM plates at 20°C. *C. elegans* were transferred daily to fresh NGM plates seeded with overnight cultures of *S. aureus* or *E. coli* OP50. Lifespan quantification performed in three independent trials; all data combined. No significant differences observed between *WT* and *glo-1* mutants under each condition. B) Defecation intervals of individual *C. elegans* observed over 10-minute time periods. *C. elegans* were transferred to fresh plates containing overnight cultures of *S. aureus* or *E. coli* OP50 and allowed to equilibrate for 1 hour. Some *C. elegans* fed *S. aureus* failed to defecate in the time allotted and were not included here; thus, statistics were not performed. However, there is a clear trend towards longer and less regular defecation intervals in both *WT* and *glo-1* mutant *C. elegans* fed *S. aureus*. C) Developmental outcomes of offspring of *C. elegans* raised on the bacteria indicated. Embryos were laid by young adult *C. elegans* exposed to the bacteria indicated from L4 stage onwards (roughly 16 hours). Embryos were laid over a period of 1-2 hours and then allowed to develop on the bacteria indicated. Counts were made twice daily for three days to observe developmental outcomes. Fisher’s Chi Square t-test. *** = p < 0.001.

## Notes

### Competing Interest Statement

The authors have declared no competing interest.

### Summary of Updates

Figure S6C added and the text is altered to reflect that addition. Some statistics have been calculated.

## REFERENCES

1. Back P, De Vos WH, Depuydt GG, Matthijssens F, Vanfleteren JR, Braeckman BP. Exploring real-time in vivo redox biology of developing and aging *Caenorhabditis elegans*. Free Radic Biol Med. 2012 Mar 1;52(5):850–9. doi: 10.1016/j.freeradbiomed.2011.11.037. Epub 2011 Dec 23. PMID: 22226831.

2. Belousov VV, Fradkov AF, Lukyanov KA, Staroverov DB, Shakhbazov KS, Terskikh AV, Lukyanov S. Genetically encoded fluorescent indicator for intracellular hydrogen peroxide. Nat Methods. 2006 Apr;3(4):281–6. doi: 10.1038/nmeth866. PMID: 16554833.

3. Bowman SL, Bi-Karchin J, Le L, Marks MS. The road to lysosome-related organelles: Insights from Hermansky-Pudlak syndrome and other rare diseases. Traffic. 2019 Jun;20(6):404–435. doi: 10.1111/tra.12646. PMID: 30945407; PMCID: PMC6541516.

4. Brenner S. The genetics of *Caenorhabditis elegans*. Genetics. 1974 May;77(1):71–94. doi: 10.1093/genetics/77.1.71. PMID: 4366476; PMCID: PMC1213120.

5. Brito S, Kim J, Bin BH. Zinc as a master regulator of intracellular organelle homeostasis. Exp Mol Med. 2026 May 1. doi: 10.1038/s12276-026-01706-2. Epub ahead of print. PMID: 42067618.

6. Chen AJ, Yuan X, Li J, Dong P, Hamza I, Cheng JX. Label-Free Imaging of Heme Dynamics in Living Organisms by Transient Absorption Microscopy. Anal Chem. 2018 Mar 6;90(5):3395–3401. doi: 10.1021/acs.analchem.7b05046. Epub 2018 Feb 14. PMID: 29401392; PMCID: PMC5972037.

7. Chen LB. Mitochondrial membrane potential in living cells. Annu Rev Cell Biol. 1988;4:155–81. doi: 10.1146/annurev.cb.04.110188.001103. PMID: 3058159.

8. Clokey GV, Jacobson LA. The autofluorescent "lipofuscin granules" in the intestinal cells of Caenorhabditis elegans are secondary lysosomes. Mech Ageing Dev. 1986 Jun;35(1):79–94. doi: 10.1016/0047-6374(86)90068-0. PMID: 3736133.

9. Coburn C, Gems D. The mysterious case of the *C. elegans* gut granule: death fluorescence, anthranilic acid and the kynurenine pathway. Front Genet. 2013 Aug 7;4:151. doi: 10.3389/fgene.2013.00151. PMID: 23967012; PMCID: PMC3735983.

10. Cogliati S, Frezza C, Soriano ME, Varanita T, Quintana-Cabrera R, Corrado M, Cipolat S, Costa V, Casarin A, Gomes LC, Perales-Clemente E, Salviati L, Fernandez-Silva P, Enriquez JA, Scorrano L. Mitochondrial cristae shape determines respiratory chain supercomplexes assembly and respiratory efficiency. Cell. 2013 Sep 26;155(1):160–71. doi: 10.1016/j.cell.2013.08.032. Epub 2013 Sep 19. PMID: 24055366; PMCID: PMC3790458.

11. Croce A, Cassata G, Disanza A, Gagliani MC, Tacchetti C, Malabarba MG, Carlier MF, Scita G, Baumeister R, Di Fiore PP. A novel actin barbed-end-capping activity in EPS-8 regulates apical morphogenesis in intestinal cells of *Caenorhabditis elegans*. Nat Cell Biol. 2004 Dec;6(12):1173–9. doi: 10.1038/ncb1198. Epub 2004 Nov 21. PMID: 15558032.

12. Cuevas-Mora K, Roque W, Shaghaghi H, Gochuico BR, Rosas IO, Summer R, Romero F. Hermansky-Pudlak syndrome-2 alters mitochondrial homeostasis in the alveolar epithelium of the lung. Respir Res. 2021 Feb 8;22(1):49. doi: 10.1186/s12931-021-01640-z. PMID: 33557836; PMCID: PMC7871590.

13. de Voer G, Peters D, Taschner PE. *Caenorhabditis elegans* as a model for lysosomal storage disorders. Biochim Biophys Acta. 2008 Jul-Aug;1782(7-8):433–46. doi: 10.1016/j.bbadis.2008.04.003. Epub 2008 May 1. PMID: 18501720.

14. Del Borrello S, Lautens M, Dolan K, Tan JH, Davie T, Schertzberg MR, Spensley MA, Caudy AA, Fraser AG. Rhodoquinone biosynthesis in *C. elegans* requires precursors generated by the kynurenine pathway. Elife. 2019 Jun 24;8:e48165. doi: 10.7554/eLife.48165. PMID: 31232688; PMCID: PMC6656428.

15. Dimov I, Maduro MF. The *C. elegans* intestine: organogenesis, digestion, and physiology. Cell Tissue Res. 2019 Sep;377(3):383–396. doi: 10.1007/s00441-019-03036-4. Epub 2019 May 7. PMID: 31065800.

16. Dooley CT, Dore TM, Hanson GT, Jackson WC, Remington SJ, Tsien RY. Imaging dynamic redox changes in mammalian cells with green fluorescent protein indicators. J Biol Chem. 2004 May 21;279(21):22284–93. doi: 10.1074/jbc.M312847200. Epub 2004 Feb 25. PMID: 14985369.

17. Estes KA, Dunbar TL, Powell JR, Ausubel FM, Troemel ER. bZIP transcription factor zip-2 mediates an early response to *Pseudomonas aeruginosa* infection in *Caenorhabditis elegans*. Proc Natl Acad Sci U S A. 2010 Feb 2;107(5):2153–8. doi: 10.1073/pnas.0914643107. Epub 2010 Jan 21. PMID: 20133860; PMCID: PMC2836710.

18. Hajdú G, Somogyvári M, Csermely P, Sőti C. Lysosome-related organelles promote stress and immune responses in *C. elegans*. Commun Biol. 2023 Sep 13;6(1):936. doi: 10.1038/s42003-023-05246-7. PMID: 37704756; PMCID: PMC10499889.

19. Hancock RE, Nijnik A, Philpott DJ. Modulating immunity as a therapy for bacterial infections. Nat Rev Microbiol. 2012 Mar 16;10(4):243–54. doi: 10.1038/nrmicro2745. PMID: 22421877.

20. Hao T, Yu J, Wu Z, Jiang J, Gong L, Wang B, Guo H, Zhao H, Lu B, Engelender S, He H, Song Z. Hypoxia-reprogramed megamitochondrion contacts and engulfs lysosome to mediate mitochondrial self-digestion. Nat Commun. 2023 Jul 11;14(1):4105. doi: 10.1038/s41467-023-39811-9. PMID: 37433770; PMCID: PMC10336010.

21. Hermann GJ, Schroeder LK, Hieb CA, Kershner AM, Rabbitts BM, Fonarev P, Grant BD, Priess JR. Genetic analysis of lysosomal trafficking in *Caenorhabditis elegans*. Mol Biol Cell. 2005 Jul;16(7):3273–88. doi: 10.1091/mbc.e05-01-0060. Epub 2005 Apr 20. PMID: 15843430; PMCID: PMC1165410.

22. Irazoqui JE, Troemel ER, Feinbaum RL, Luhachack LG, Cezairliyan BO, Ausubel FM. Distinct pathogenesis and host responses during infection of *C. elegans* by *P. aeruginosa* and *S. aureus*. PLoS Pathog. 2010 Jul 1;6(7):e1000982. doi: 10.1371/journal.ppat.1000982. PMID: 20617181; PMCID: PMC2895663.

23. Kapetanovic R, Afroz SF, Curson JEB, Kirmes I, Ramnath D, Nothjunge S, Liu J, Atkinson JD, Ahier A, Raven KD, Bosch M, Keller B, Lawrence GMEP, Gupta KD, Shakespear MR, Ferguson C, Stocks CJ, Bokil NJ, Matthias G, Nguyen TTK, Khalil ZG, Reid RC, Hansford KA, Hansbro PM, Cooper MA, Schembri MA, Blumenthal A, Schroder K, Fairlie DP, Pol A, Matthias P, Parton RG, Zuryn S, Sweet MJ. Mitochondrial fission mediates an evolutionarily conserved antibacterial defense response. Sci Immunol. 2026 Apr 24;11(118):eaed2623. doi: 10.1126/sciimmunol.aed2623. Epub 2026 Apr 24. PMID: 42030373.

24. Kirienko NV, Cezairliyan BO, Ausubel FM, Powell JR. *Pseudomonas aeruginosa* PA14 pathogenesis in *Caenorhabditis elegans*. Methods Mol Biol. 2014;1149:653–69. doi: 10.1007/978-1-4939-0473-0_50. PMID: 24818940.

25. Kook S, Wang P, Young LR, Schwake M, Saftig P, Weng X, Meng Y, Neculai D, Marks MS, Gonzales L, Beers MF, Guttentag S. Impaired Lysosomal Integral Membrane Protein 2-dependent Peroxiredoxin 6 Delivery to Lamellar Bodies Accounts for Altered Alveolar Phospholipid Content in Adaptor Protein-3-deficient pearl Mice. J Biol Chem. 2016 Apr 15;291(16):8414–27. doi: 10.1074/jbc.M116.720201. Epub 2016 Feb 23. PMID: 26907692; PMCID: PMC4861416.

26. Labbé RF, Vreman HJ, Stevenson DK. Zinc protoporphyrin: A metabolite with a mission. Clin Chem. 1999 Dec;45(12):2060–72. PMID: 10585337.

27. Leiser SF, Miller H, Rossner R, Fletcher M, Leonard A, Primitivo M, Rintala N, Ramos FJ, Miller DL, Kaeberlein M. Cell nonautonomous activation of flavin-containing monooxygenase promotes longevity and health span. Science. 2015 Dec 11;350(6266):1375–1378. doi: 10.1126/science.aac9257. Epub 2015 Nov 19. PMID: 26586189; PMCID: PMC4801033.

28. Li X, Straub J, Medeiros TC, Mehra C, den Brave F, Peker E, Atanassov I, Stillger K, Michaelis JB, Burbridge E, Adrain C, Münch C, Riemer J, Becker T, Pernas LF. Mitochondria shed their outer membrane in response to infection-induced stress. Science. 2022 Jan 14;375(6577):eabi4343. doi: 10.1126/science.abi4343. Epub 2022 Jan 14. PMID: 35025629.

29. Liberati NT, Fitzgerald KA, Kim DH, Feinbaum R, Golenbock DT, Ausubel FM. Requirement for a conserved Toll/interleukin-1 resistance domain protein in the *Caenorhabditis elegans* immune response. Proc Natl Acad Sci U S A. 2004 Apr 27;101(17):6593–8. doi: 10.1073/pnas.0308625101. PMID: 15123841; PMCID: PMC404090.

30. Liu Q, Yoo S, Zhang ZA, Li L, Su H, Vannur L, Wooldredge AC, Hughes JB, Desprez PY, Hao N, Lithgow G, Andersen JK, Hansen M, Campisi J, Zhou C. Mitochondria-lysosome coupling contributes to lysosome acidification and aging. Mol Cell. 2026 Jun 18;86(12):2425–2442.e10. doi: 10.1016/j.molcel.2026.05.004. Epub 2026 May 29. PMID: 42214330; PMCID: PMC13263124.

31. Liu Y, Samuel BS, Breen PC, Ruvkun G. *Caenorhabditis elegans* pathways that surveil and defend mitochondria. Nature. 2014 Apr 17;508(7496):406–10. doi: 10.1038/nature13204. Epub 2014 Apr 2. PMID: 24695221; PMCID: PMC4102179.

32. Mallick A, Haynes CM. Methods to analyze the mitochondrial unfolded protein response (UPRmt). Methods Enzymol. 2024;707:543–564. doi: 10.1016/bs.mie.2024.07.029. Epub 2024 Aug 12. PMID: 39488390.

33. Marshall JH, Wilmoth GJ. Pigments of *Staphylococcus aureus*, a series of triterpenoid carotenoids. J Bacteriol. 1981 Sep;147(3):900–13. doi: 10.1128/jb.147.3.900-913.1981. PMID: 7275936; PMCID: PMC216126.

34. Mao K, Ji F, Breen P, Sewell A, Han M, Sadreyev R, Ruvkun G. Mitochondrial Dysfunction in *C. elegans* Activates Mitochondrial Relocalization and Nuclear Hormone Receptor-Dependent Detoxification Genes. Cell Metab. 2019 May 7;29(5):1182–1191.e4. doi: 10.1016/j.cmet.2019.01.022. Epub 2019 Feb 21. PMID: 30799287; PMCID: PMC6506380.

35. McEwan DL, Kirienko NV, Ausubel FM. Host translational inhibition by *Pseudomonas aeruginosa* Exotoxin A Triggers an immune response in *Caenorhabditis elegans*. Cell Host Microbe. 2012 Apr 19;11(4):364–74. doi: 10.1016/j.chom.2012.02.007. PMID: 22520464; PMCID: PMC3334877.

36. Meng R, Wu J, Harper DC, Wang Y, Kowalska MA, Abrams CS, Brass LF, Poncz M, Stalker TJ, Marks MS. Defective release of α granule and lysosome contents from platelets in mouse Hermansky-Pudlak syndrome models. Blood. 2015 Mar 5;125(10):1623–32. doi: 10.1182/blood-2014-07-586727. Epub 2014 Dec 4. PMID: 25477496; PMCID: PMC4351507.

37. Morris C, Foster OK, Handa S, Peloza K, Voss L, Somhegyi H, Jian Y, Vo MV, Harp M, Rambo FM, Yang C, Hermann GJ. Function and regulation of the *Caenorhabditis elegans* Rab32 family member *GLO-1* in lysosome-related organelle biogenesis. PLoS Genet. 2018 Nov 12;14(11):e1007772. doi: 10.1371/journal.pgen.1007772. PMID: 30419011; PMCID: PMC6268011.

38. O’Neill AJ. *Staphylococcus aureus* SH1000 and 8325-4: comparative genome sequences of key laboratory strains in staphylococcal research. Lett Appl Microbiol. 2010 Sep;51(3):358–61. doi: 10.1111/j.1472-765X.2010.02885.x. Epub 2010 Jun 10. PMID: 20618890.

39. O’Rourke EJ, Ruvkun G. MXL-3 and HLH-30 transcriptionally link lipolysis and autophagy to nutrient availability. Nat Cell Biol. 2013 Jun;15(6):668–76. doi: 10.1038/ncb2741. Epub 2013 Apr 21. Erratum in: Nat Cell Biol. 2015 Jan;17(1):104. PMID: 23604316; PMCID: PMC3723461.

40. Pukkila-Worley R, Ausubel FM. Immune defense mechanisms in the *Caenorhabditis elegans* intestinal epithelium. Curr Opin Immunol. 2012 Feb;24(1):3–9. doi: 10.1016/j.coi.2011.10.004. Epub 2012 Jan 9. PMID: 22236697; PMCID: PMC3660727.

41. Poulos TL. Heme enzyme structure and function. Chem Rev. 2014 Apr 9;114(7):3919–62. doi: 10.1021/cr400415k. Epub 2014 Jan 8. PMID: 24400737; PMCID: PMC3981943.

42. Rae R, Witte H, Rödelsperger C, Sommer RJ. The importance of being regular: *Caenorhabditis elegans* and *Pristionchus pacificus* defecation mutants are hypersusceptible to bacterial pathogens. Int J Parasitol. 2012 Jul;42(8):747–53. doi: 10.1016/j.ijpara.2012.05.005. Epub 2012 Jun 13. PMID: 22705203.

43. Roberts Buceta PM, Romanelli-Cedrez L, Babcock SJ, Xun H, VonPaige ML, Higley TW, Schlatter TD, Davis DC, Drexelius JA, Culver JC, Carrera I, Shepherd JN, Salinas G. The kynurenine pathway is essential for rhodoquinone biosynthesis in *Caenorhabditis elegans*. J Biol Chem. 2019 Jul 12;294(28):11047–11053. doi: 10.1074/jbc.AC119.009475. Epub 2019 Jun 7. PMID: 31177094; PMCID: PMC6635453.

44. Roh HC, Collier S, Guthrie J, Robertson JD, Kornfeld K. Lysosome-related organelles in intestinal cells are a zinc storage site in *C. elegans*. Cell Metab. 2012 Jan 4;15(1):88–99. doi: 10.1016/j.cmet.2011.12.003. PMID: 22225878; PMCID: PMC4026189.

45. Salinas G, Langelaan DN, Shepherd JN. Rhodoquinone in bacteria and animals: Two distinct pathways for biosynthesis of this key electron transporter used in anaerobic bioenergetics. Biochim Biophys Acta Bioenerg. 2020 Nov 1;1861(11):148278. doi: 10.1016/j.bbabio.2020.148278. Epub 2020 Jul 28. PMID: 32735860.

46. Schroeder LK, Kremer S, Kramer MJ, Currie E, Kwan E, Watts JL, Lawrenson AL, Hermann GJ. Function of the *Caenorhabditis elegans* ABC transporter PGP-2 in the biogenesis of a lysosome-related fat storage organelle. Mol Biol Cell. 2007 Mar;18(3):995–1008. doi: 10.1091/mbc.e06-08-0685. Epub 2007 Jan 3. PMID: 17202409; PMCID: PMC1805080.

47. Sifri CD, Begun J, Ausubel FM, Calderwood SB. *Caenorhabditis elegans* as a model host for *Staphylococcus aureus* pathogenesis. Infect Immun. 2003 Apr;71(4):2208–17. doi: 10.1128/IAI.71.4.2208-2217.2003. PMID: 12654843; PMCID: PMC152095.

48. Tan CH, Wang TY, Park H, Lomenick B, Chou TF, Sternberg PW. Single-tissue proteomics in *Caenorhabditis elegans* reveals proteins resident in intestinal lysosome-related organelles. Proc Natl Acad Sci U S A. 2024 Jun 18;121(25):e2322588121. doi: 10.1073/pnas.2322588121. Epub 2024 Jun 11. PMID: 38861598; PMCID: PMC11194598.

49. Tiku V, Tan MW, Dikic I. Mitochondrial Functions in Infection and Immunity. Trends Cell Biol. 2020 Apr;30(4):263–275. doi: 10.1016/j.tcb.2020.01.006. Epub 2020 Feb 11. Erratum in: Trends Cell Biol. 2020 Sep;30(9):748. doi: 10.1016/j.tcb.2020.07.001. PMID: 32200805; PMCID: PMC7126537.

50. Tse-Kang S, Wani KA, Pukkila-Worley R. Patterns of pathogenesis in innate immunity: insights from *C. elegans*. Nat Rev Immunol. 2025 Sep;25(9):637–648. doi: 10.1038/s41577-025-01167-0. Epub 2025 Apr 17. PMID: 40247006; PMCID: PMC13034488.

51. Tse-Kang SY, Wani KA, Peterson ND, Page A, Humphries F, Pukkila-Worley R. Intestinal immunity in *C. elegans* is activated by pathogen effector-triggered aggregation of the guard protein TIR-1 on lysosome-related organelles. Immunity. 2024 Oct 8;57(10):2280–2295.e6. doi: 10.1016/j.immuni.2024.08.013. Epub 2024 Sep 18. PMID: 39299238; PMCID: PMC11464196.

52. Valera-Alberni M, Yao P, Romero-Sanz S, Lanjuin A, Mair WB. Novel Imaging Tools to Study Mitochondrial Dynamics in *Caenorhabditis elegans*. bioRxiv [Preprint]. 2024 Jul 16:2024.07.16.603730. doi: 10.1101/2024.07.16.603730. Update in: Life Sci Alliance. 2024 Sep 11;7(11):e202402918. doi: 10.26508/lsa.202402918. PMID: 39071403; PMCID: PMC11275731.

53. Visvikis O, Ihuegbu N, Labed SA, Luhachack LG, Alves AF, Wollenberg AC, Stuart LM, Stormo GD, Irazoqui JE. Innate host defense requires TFEB-mediated transcription of cytoprotective and antimicrobial genes. Immunity. 2014 Jun 19;40(6):896–909. doi: 10.1016/j.immuni.2014.05.002. Epub 2014 May 29. PMID: 24882217; PMCID: PMC4104614.

54. Wang JY, Young LR. Hermansky-Pudlak Syndrome. Clin Chest Med. 2025 Dec;46(4):701–710. doi: 10.1016/j.ccm.2025.07.009. Epub 2025 Sep 1. PMID: 41110930.

55. Weimer RM. Preservation of *C. elegans* tissue via high-pressure freezing and freeze-substitution for ultrastructural analysis and immunocytochemistry. Methods Mol Biol. 2006;351:203–21. doi: 10.1385/1-59745-151-7:203. PMID: 16988436.

56. Yoneda T, Benedetti C, Urano F, Clark SG, Harding HP, Ron D. Compartment-specific perturbation of protein handling activates genes encoding mitochondrial chaperones. J Cell Sci. 2004 Aug 15;117(Pt 18):4055–66. doi: 10.1242/jcs.01275. Epub 2004 Jul 27. PMID: 15280428.

57. Zhang SO, Box AC, Xu N, Le Men J, Yu J, Guo F, Trimble R, Mak HY. Genetic and dietary regulation of lipid droplet expansion in *Caenorhabditis elegans*. Proc Natl Acad Sci U S A. 2010 Mar 9;107(10):4640–5. doi: 10.1073/pnas.0912308107. Epub 2010 Feb 22. PMID: 20176933; PMCID: PMC2842062.

58. Zhang P, Medwig-Kinney TN, Goldstein B. Architecture of the cortical actomyosin network driving apical constriction in *C. elegans*. J Cell Biol. 2023 Sep 4;222(9):e202302102. doi: 10.1083/jcb.202302102. Epub 2023 Jun 23. PMID: 37351566; PMCID: PMC10289891.

59. Zhou X, Liu K, Li J, Cui L, Dong J, Li J, Meng X, Zhu G, Wang H. PINK1/Parkin-mediated mitophagy enhances the survival of *Staphylococcus aureus* in bovine macrophages. J Cell Mol Med. 2023 Feb;27(3):412–421. doi: 10.1111/jcmm.17664. Epub 2023 Jan 9. PMID: 36625039; PMCID: PMC9889626.

