## Supplemental Table 1 for "Lysosome-related organelle genes are required for mitochondrial transformations during bacterial infections in *Caenorhabditis elegans*"

Table S1. Strains used in this study

| **Strain name** | **Genotype** | **Source** |
|---|---|---|
| N2 | *Wild type* | |
| JJ1271 | *glo-1(zu391) X* | Hermann et al., 2005 |
| JV1 | *unc-119(ed3) III; jrIs1[rpl-17p::HyPer + unc-119(+)]* | Back et al., 2012 |
| JV2 | *unc-119(ed3) III;<br>jrIs2 [rpl-17p::Grx1-roGFP2 + unc-119(+)]* | Back et al., 2012 |
| KAE28 | *unc-119(ed3) III;<br>seaEx13[fmo-2p::GFP + unc-119(+)]* | Leiser et al., 2015 |
| LP898 | *eps-8(cp441[eps-8::mNG-C1]) IV* | Zhang et al., 2023 |
| SJ4100 | *zcIs13 [hsp-6p::GFP + lin-15(+)]* | Yoneda et al., 2004 |
| WBM1689 | *tomm-70(wbm81[tomm-70::GFP) III;<br>scpl-4(wbm108[scpl-4::wrmScarlet]) V* | Valera-Alberni et al., 2024 |
| GR2250 | *mgIs73[cyp-14A4p::GFP::cyp-14A4 3'UTR + myo-3p::mCherry::unc-54 3'UTR] V* | Mao et al., 2019 |
| GR2267 | *mgTi33[vha-6p::tomm-20(1-54)::mScarlet::nhr-45 3'UTR]* | Mao et al., 2019 |
| GR3713 | *glo-4(ok623) V; mgTi33[vha-6p::tomm-20(1-54) ::mScarlet::nhr-45 3'UTR]* | This study |
| GR3714 | *mgTi33[vha-6p::tomm-20(1-54)::mScarlet::nhr-45 3'UTR]; agIs17[irg-1::GFP]* | This study |
| GR3716 | *mgTi33[vha-6p::tomm-20(1-54)::mScarlet::nhr-45 3'UTR];<br>glo-1(zu391) X; kxEx15[ges-1pro::GLO-1::GFP]* | This study |
| GR3722 | *mgTi33[vha-6p::tomm-20(1-54)::mScarlet::nhr-45 3'UTR]; agIs17[irg-1::GFP]; glo-1(zu391) X* | This study |
| GR3725 | *glo-4(ok623) V;<br>seaEx13 [fmo-2p::GFP + unc-119(+)]* | This study |
| GR3733 | *glo-1(zu391) X;<br>seaEx13 [fmo-2p::GFP + unc-119(+)]* | This study |
| GR3765 | *tomm-70(wbm81[tomm-70::GFP]) III; scpl-4(wbm108[scpl-4::wrmScarlet]) V; glo-1(zu391) X* | This study |
| GR3766 | *pgp-2(kx48) I; mgTi33[vha-6p::tomm-20(1-54)::mScarlet::nhr-45 3'UTR]; agIs17[irg-1::GFP]* | This study |
| GR3768 | *pgp-2(kx48) I; seaEx13 [fmo-2p::GFP + unc-119(+)]* | This study |
| GR3770 | *glo-4(ok623) V; mgTi33[vha-6p::tomm-20(1-54)::mScarlet::nhr-45 3'UTR]; agIs17[irg-1::GFP]* | This study |
| GR3772 | *pgp-2(kx48) I; tomm-70(wbm81[tomm-70::GFP]) III; scpl-4(wbm108[scpl-4::wrmScarlet]) V* | This study |
| GR3774 | *pgp-2(kx48) I; mgTi33[vha-6p::tomm-20(1-54)::mScarlet::nhr-45 3'UTR]* | This study |
| GR3791 | *mgTi33[vha-6p::tomm-20(1-54)::mScarlet::nhr-45 3'UTR]; glo-1(zu391) X* | This study |
